# Immune, Autonomic, and Endocrine Dysregulation in Autism and Ehlers-Danlos Syndrome/Hypermobility Spectrum Disorders Versus Unaffected Controls

**DOI:** 10.1101/670661

**Authors:** Emily L. Casanova, Julia L. Sharp, Stephen M. Edelson, Desmond P. Kelly, Estate M. Sokhadze, Manuel F. Casanova

## Abstract

**Background:** A growing body of literature suggests etiological overlap between Ehlers-Danlos syndrome (EDS)/hypermobility spectrum disorders (HSD) and some cases of autism, although this relationship is poorly delineated. In addition, immune, autonomic, and endocrine dysregulation are reported in both conditions and may be relevant to their respective etiologies.

**Aims:** To study symptom overlap in these two comorbid spectrum conditions.

**Methods and Procedures:** We surveyed 702 adults aged 25+ years on a variety of EDS/HSD-related health topics, comparing individuals with EDS/HSD, autism, and unaffected controls.

**Outcomes and Results:** The autism group reported similar though less severe symptomology as the EDS/HSD group, especially in areas of immune/autonomic/endocrine dysregulation, connective tissue abnormalities (i.e., skin, bruising/bleeding), and chronic pain. EDS/HSD mothers with autistic children reported more immune symptoms than EDS/HSD mothers without, suggesting the maternal immune system could play a heritable role in these conditions (*p* = 0.0119).

**Conclusions and Implications:** These data suggest that EDS/HSD and autism share aspects of immune/autonomic/endocrine dysregulation, pain, and some tissue fragility, which is typically more severe in the former. This overlap, as well as documented comorbidity, suggests some forms of autism may be hereditary connective tissue disorders (HCTD).

## INTRODUCTION

At face value, Ehlers-Danlos syndrome (EDS)/hypermobility spectrum disorders (HSD) and autism appear to have little in common. Though autism can be accompanied by a variety of medical issues, it is first and foremost defined as a behavioral syndrome. In contrast, EDS/HSD is a group of hereditary connective tissue disorders (HCTD) believed to be due to a deficiency in collagen fibrillogenesis. Therefore, symptoms in these conditions stem largely from connective tissue dysplasias (joint instability, pain, skin elasticity, tissue fragility, etc.).

Early evidence suggests that autism and EDS/HSD overlap comorbidly (Casanova et al., 2018; Baeza-Velasco et al., 2015). Although a number of case studies have been published since the 1980s, a more recent nationwide study from Sweden indicates that approximately 3% of individuals with EDS have an autism diagnosis (Fehlow & Tennstedt, 1985; Sieg, 1992; Fehlow et al., 1993; Takei et al., 2011; Cederlöf et al., 2016). However, the high preponderance of women with EDS/HSD and the recognition that autism is often under-diagnosed in female demographics suggests this could be an underestimate of comorbidity (Gould & Ashton-Smith, 2011). Up-to-date prevalence estimates of hypermobile EDS (hEDS), the most common form of EDS, are also currently lacking, as are estimates on the new diagnostic entity, hypermobility spectrum disorders (HSD).

Although etiological mechanisms underlying the co-occurrence of EDS/HSD and autism are poorly understood, we know for instance that the extracellular matrix (ECM) plays foundational roles in brain development, providing scaffolds for proliferating and migrating neurons, as well as helping to maintain synaptic contacts throughout life (Mercier et al., 2002; Thomas & Steindler, 1996; Sheen et al., 2005; Su et al., 2010). The immune system has also been implicated in autism’s etiology and is a system that is significantly dysregulated in EDS/HSD and other hereditary connective tissue disorders (Seneviratne, 2017; Felgentreff et al., 2014; Patterson, 2011). In addition, autonomic disorders have been reported in both these groups and could be a cause or consequence of immune dysregulation (Sokhadze et al., 2018; Cheung & Vadas, 2015; Czura & Tracey, 2005; Roth et al., 2004; Bienenstock et al., 1987). For these reasons, we have elected to study self-reported immune and autonomic symptom frequency in the adult EDS/HSD and autism populations as compared to controls. We also investigate the frequency in autism of other symptoms typically associated with EDS/HSD, including items such as chronic pain and tissue fragility.

## MATERIAL AND METHODS

### Study Population

Approximately 99% of respondents answered the survey on behalf of themselves, rather than as legal guardians of adult wards. Our study sample was composed of English-speaking adults aged 25 years or older [*N* = 702]. (For basic demographics, see Table 1. Note that specific data on socioeconomic status and educational attainment levels were not recorded.)

**Table 1.** Sex, age, racial descent, and country of residence according to group. Figures presented represent diagnostically “pure” (i.e., non-mixed) clinical and control groups.

| Table 1. Participant Demographics |  |  |  |  |
| --- | --- | --- | --- | --- |
| Demographics |  | EDS/HSD<br>(N = 268) | Autism<br>(N = 152) | CON<br>(N = 147) |
| *Sex* | Male | 6% | 34.9% | 31.8% |
|  | Female | 92.5% | 58.6% | 66.2% |
|  | Other | 1.5% | 6.6% | 2% |
| *Age* | Average | 35.9 ± 9.7<br>years | 37.6 ± 10.4<br>years | 33.0 ± 10.2<br>years |
| *Descent* | Sub-saharan African | 2.6% | 2% | 0.7% |
|  | Middle Eastern & Northern African | 1.5% | 0.7% | 2% |
|  | East Asian & Native American | 13.8% | 11.1% | 8.1% |
|  | South Asian | 1.1% | 0% | 0% |
|  | Northwestern European | 85.4% | 87% | 78.4% |
|  | Eastern European | 23.5% | 17.6% | 27% |
|  | Southern European | 17.5% | 8.5% | 17.6% |
| *Country of Residence* | Australia/New Zealand | 4.9% | 8.6% | 2% |
|  | Bahrain | 0% | 0% | 0.7% |
|  | Belgium | 0.4% | 1.3% | 0% |
|  | Brazil | 0% | 0% | 0.7% |
|  | Canada | 9.7% | 5.3% | 10.1% |
|  | Democratic Republic of Congo | 0% | 0% | 0.7% |
|  | Denmark | 0.7% | 1.3% | 0% |
|  | Dominican Republic | 0% | 0% | 0.7% |
|  | Finland | 0% | 0% | 0.7% |
|  | France | 0% | 3.3% | 0% |
|  | Germany | 0.4% | 0.7% | 2.7% |
|  | Iceland | 0% | 2% | 0% |
|  | Israel | 0% | 0% | 0.7% |
|  | Italy | 0% | 0% | 0.7% |
|  | Netherlands | 0% | 2% | 0% |
|  | Norway | 0% | 0.7% | 0.7% |
|  | Poland | 0.4% | 0% | 1.4% |
|  | Russia | 0.4% | 0.7% | 1.4% |
|  | Singapore | 0.4% | 0% | 0% |
|  | South Africa | 0.4% | 0.7% | 0.7% |
|  | Spain | 0% | 0.7% | 0% |
|  | Sweden | 0.4% | 1.3% | 0% |
|  | United Kingdom | 9.3% | 25% | 11.5% |
|  | United States of America | 71.3% | 45.4% | 61.5% |
|  | Uzbekistan | 0.4% | 0% | 0.7% |

Our sample was divided into three major groups: 1) individuals reporting diagnosis of EDS or the now-obsolete joint hypermobility syndrome (JHS), with or without an additional diagnosis of autism [*N* = 403]; 2) individuals reporting a diagnosis of autism but without a diagnosis of EDS/HSD [*N* = 152]; and 3) individuals reporting neither EDS/HSD or autism (diagnosed or suspected) or with a 1^st^ degree relative with any of these diagnoses [*N* = 147]. Individuals who were under the age of 25 or who had a 1^st^ degree relative with EDS/HSD or autism and didn’t themselves have either of these conditions were removed from the study. (Unfortunately, family members of those with EDS/HSD or autism did not form large enough subgroups to be studied individually.) In addition, individuals who suspected a diagnosis of EDS/HSD but did not report a diagnosis were also removed from the study. For most analyses, the data of individuals with EDS/HSD diagnoses who suspected they had autism but were undiagnosed were not used, with a few exceptions such as the analyses on maternal heredity and EDS/HSD within-group comparisons. For analyses in which sex was controlled as a potential confounding variable, individuals who did not identify as either male or female were removed from the data pool.

The majority (74%) of EDS/HSD respondents reported a diagnosis of hEDS [*N* = 294]. Meanwhile, 9% of EDS/HSD respondents reported a diagnosis of classic EDS (cEDS) [*N* = 37], 3% reported a diagnosis of vascular EDS (vEDS) [*N* = 12], and 6% reported another form of EDS not easily categorized in this study. Nine percent of EDS/HSD respondents reported the older diagnosis of JHS, which would nowadays either be diagnosed as generalized hypermobility spectrum disorder (G-HSD) or asymptomatic generalized joint hypermobility (A-GJH). In this case, however, all JHS respondents reported significant musculoskeletal pain and instability, indicating that they are symptomatic and would therefore fall under the G-HSD umbrella were they to be reassessed at present.

### Survey

This survey was approved by the Institutional Review Board (IRB) of the Greenville Health System (GHS) (ID: Pro00073030). It was an extension of a previous survey designed by our group, partly based off of the *Vanderbilt Autonomic Dysfunction Center Medical Questionnaire* and a series of surveys utilized by the Autism Research Institute (see Casanova et al., 2018; Vanderbilt Autonomic Dysfunction Center, 2018).

The survey was built on and hosted by the website, SurveyGizmo, and was advertised on a variety of websites, including Ehlers-Danlos-specific Facebook and reddit groups, Wrong Planet forums, and Survey Tandem. For websites such as reddit and Survey Tandem, requirements for posting research advertisements were followed according to the rules of each respective site (see Supplementary File 1 for survey). For forums and Facebook groups, the administrative teams of these webgroups were approached and asked for permission in order to post the study advertisement. Following administrator approval, the primary author posted the study advert.

The survey weblink (http://www.surveygizmo.com/s3/3420093/Expanding-Our-Understanding-of-Symptoms-Associated-with-Ehlers-Danlos-Syndromes) led to a description of the study, including potential risks/benefits and expectations, investigator contact information, and a waiver of consent. The survey was available and data was collected for six months.

Survey questions covered a wide variety of medical and health issues, many of which have been referenced in the EDS-related literature as well as our own previous work (Casanova et al., 2018). Some of the major systems of interest include the immune (Fig. 1a), endocrine, and autonomic nervous systems, which have previously been implicated in both EDS/HSD and autism (Seneviratne et al., 2017; Kushki et al., 2013; De Wandele et al., 2014; Careaga et al., 2017; Ingudomnukul et al., 2007; Castori et al., 2012). In addition, we also asked about other symptoms associated with EDS/HSD, such as musculoskeletal pain and instability, dermatological issues (elasticity, scarring, wound healing), vascular complications, and other forms of chronic pain. Some psychiatric comorbidities were also covered in the survey.

**Figure 1.**
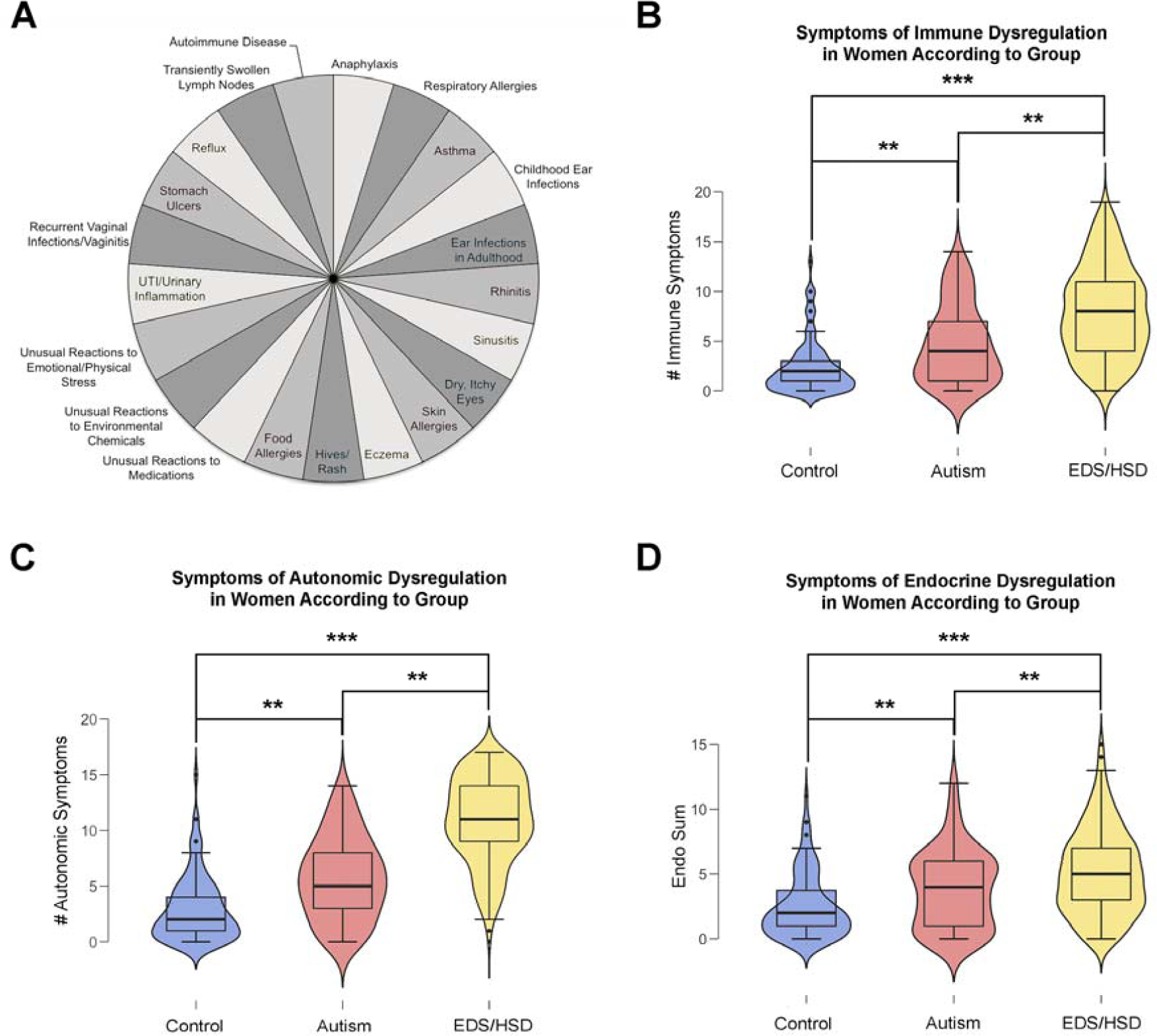
Immune, endocrine, and autonomic dysregulation. (A) Pie chart listing the various immune-mediated symptoms investigated in this study. (Slices are not proportional to rate of occurrence in this study.) (B-D) Immune-, autonomic-, and endocrine-mediated symptoms in women according to group.

### Statistical Analyses

Group comparisons for binary outcomes were compared using logistic regression analysis. When a group proportion for a particular symptom was zero or very small, odds ratio confidence intervals including that group were unbounded (upper bound was infinity) and were not interpreted. Multiple comparisons within a particular symptom were adjusted using a false discovery rate *p*-value adjustment. For comparing symptom sums among the groups, a Kruskal-Wallis test was used. Follow-up analyses to compare groups pairwise were considered using the Wilcoxon Rank Sum Test. A significance level of 0.05 was used for all tests of significance.

## RESULTS

### The Immune, Endocrine, & Autonomic Nervous Systems

In agreement with the general literature, we found that the distributions of immune-mediated and autonomic symptoms differed across our control group by gender (women > men) [*W* = 2826.5-2843, *p* = 0.021-0.024] (Oertelt, 2012; Lambrecht et al., 2010). A similar gender disparity was also present within each of our clinical groups, although the number of men within the EDS/HSD group (*N* = 16) was comparatively small given the extreme sex-skewing of the condition [*W* = 2826.5-3018, *p* < 0.001-0.016] (Hamonet & Brock, 2015). Using a series of Kruskal-Wallis rank sum tests, we found that immune-mediated and autonomic symptoms were also staggered according to group: individuals with EDS/HSD reported the most symptoms, followed by those with autism, and finally controls [*X*^*2*^ = 11.782-217.57, *p* < 0.0001-0.003] (Fig. 1b,c). In contrast, endocrine-mediated symptoms primarily differed in females by group (EDS/HSD > autism > controls) [*X*^*2*^ = 55.792, *p* < 0.0001] (Fig. 1d) but did not differ significantly in males [*X*^*2*^ = 1.6871, *p* = 0.4302]. This was due to the fact that female-specific endocrine disorders (e.g., endometriosis, menorrhagia) were the major drivers of group divergence in this study. (See Supplementary File1, Table 1 for full results.)

When we looked at EDS/HSD subtypes, we found that their immune, autonomic, and endocrine profiles were more similar than not. While we were unable to investigate all EDS subtypes due to their rarity, we were nevertheless able to compare hEDS, cEDS, vEDS, and JHS (the latter a diagnosis which many respondents retain due to the recency of nosological changes) (Castori et al., 2017). There was a significant difference in the distribution of both immune-mediated and autonomic symptoms according to EDS/HSD subtype [*X*^*2*^ = 13.205-16.234, *p* = 0.0027-0.0103], though endocrine symptoms did not appear to differ [*X*^*2*^ = 4.9388, *p* = 0.2936]. Immunologically and autonomically, JHS is less severe than the other EDS subtypes; however, symptomologically JHS is more similar to EDS than to either the autism or control groups, suggesting a difference in severity rather than necessarily one of kind (Supplementary File 2, Table 2).

**Table 2.** EDS-associated symptom frequency by group

| **Table 2. Ehlers-Danlos-related Symptoms** |  |  |  |  |
| --- | --- | --- | --- | --- |
|  | **EDS/HSD** | **Autism** | **CON** | **Respondents used in analysis** |
| *Head Injury* | 20% | 8% | 9% | females only |
| *Migraine* | 80% | 52% | 42% | females only |
| *Non-migraine headache* | 47% | 11% | 15% | females only |
| *Bruise Easily* | 87% | 46% | 32% | females only |
| *Difficult-to-control bleeds* | 29% | 1% | 2% | females only |
| *Scars easily* | 70% | 27% | 14% | females only |
| *Cigarette paper scars* | 46% | 1% | 0% | females only |
| *Elastic skin* | 47% | 6% | 0% | females only |
| *Slow wound healing* | 66% | 19% | 5% | females only |
| *Clumsy* | 66% | 52% | 19% | females only |
| *Joint pain* | 95% | 26% | 16% | females only |
| *Muscle pain* | 82% | 21% | 7% | females only |
| *Arthritis* | 29% | 12% | 6% | females only |
| *Fibromyalgia* | 29% | 10% | 2% | females only |
| *Temporomandibular joint (TMJ) pain* | 64% | 24% | 9% | females only |
| *Neuropathy* | 36% | 7% | 4% | females only |
| *Joint dislocations/subluxations* | 89% | 8% | 3% | females only |
| *Foot problems* | 90% | 42% | 33% | females only |
| *Crooked/crowded teeth (current or historical)* | 78% | 69% | 53% | females only |
| *Scoliosis* | 36% | 15% | 11% | females only |
| *Kyphosis* | 13% | 2% | 2% | females only |
| *Missing/extra bones* | 17% | 8% | 5% | females only |
| *Chiari malformation* | 6% | 0% | 0% | females only |
| *Prematurity (self)* | 13% | 16% | 12% | females only |
| *Prematurity (child)* | 25% | 9% | 24% | mothers only |
| *Preeclampsia* | 26% | 9% | 26% | mothers only |
| *Gestational diabetes* | 11% | 6% | 18% | mothers only |
| *Uterine prolapse* | 13% | 3% | 0% | mothers only |
| *Mitral valve prolapse* | 14% | 3% | 2% | females only |
| *Venous arterial dissection* | 1% | 0% | 0% | females only |
| *Cavernous sinus fistula* | 0% | 0% | 0% | females only |
| *Aneurysm* | 2% | 0% | 1% | females only |
| *Brain hemorrhage* | 0% | 1% | 0% | females only |
| *Stroke* | 1% | 1% | 0% | females only |
| *Other vascular complications* | 15% | 0% | 3% | females only |
| *Myopia* | 61% | 62% | 45% | females only |
| *Eye strain* | 52% | 26% | 5% | females only |
| *Retinal detachment* | 3% | 1% | 0% | females only |
| *Ptosis* | 21% | 4% | 2% | females only |
| *Blue/gray sclera* | 24% | 1% | 0% | females only |
| *Astigmatism* | 64% | 54% | 37% | females only |
| *Keratoconus* | 3% | 1% | 0% | females only |
| *Chronic dry eye* | 36% | 9% | 10% | females only |
| *Strabismus* | 11% | 7% | 4% | females only |
| *All cancer (self)* | 5% | 4% | 2% | females only |
| *Cancer under 35* | 4% | 0% | 1% | females only |
| *Epilepsy (self)* | 4% | 6% | 0% | females only |
| *Epilepsy (relatives)* | 8% | 5% | 1% | males and females |
| *ADHD* | 20% | 19% | 7% | females only |
| *Anxiety disorder* | 42% | 42% | 26% | females only |
| *Mood disorder* | 29% | 47% | 20% | females only |
| *PTSD* | 13% | 8% | 3% | females only |
| *Adverse reaction to vaccination* | 28% | 18% | 6% | females only |
| *Mild immunodeficiency* | 24% | 9% | 8% | females only |

There is a small but growing literature that suggests an etiological overlap between EDS/HSD and autism (Casanova et al., 2018; Baeza-Velasco et al., 2015). While we are unable to estimate true comorbidity rates in this study due to the ways in which respondents were recruited (potential bias), we will nevertheless briefly describe our EDS/HSD sample. Approximately 12% of EDS/HSD participants reported a dual diagnosis of autism. Meanwhile, an additional 21% of EDS/HSD respondents reported that they suspected they had autism or fell within the broader phenotype. Therefore, up to one-third of our EDS/HSD sample may have been on or near the autism spectrum.

We also found that a large percentage of EDS/HSD mothers reported having autistic children (20%), a figure that was almost identical to that seen in mothers with autism (19%) (Fig. 2a). Although the log odds did not significantly differ between the two groups [*X*^*2*^ = 1.1322, *p* = 0.5677], these results must be interpreted with extreme caution due to the potentially biased manner in which respondents were recruited. Interestingly, rates of autism in the offspring did not significantly differ across the different maternal EDS/HSD subgroups, which tentatively suggests that autism may be associated with various forms of EDS/HSD [*X*^*2*^ = 0.4807, *p* = 0.4881] (Fig. 2b). Further research is desperately needed to determine both autism comorbidity and inheritance rates in these families.

**Figure 2.**
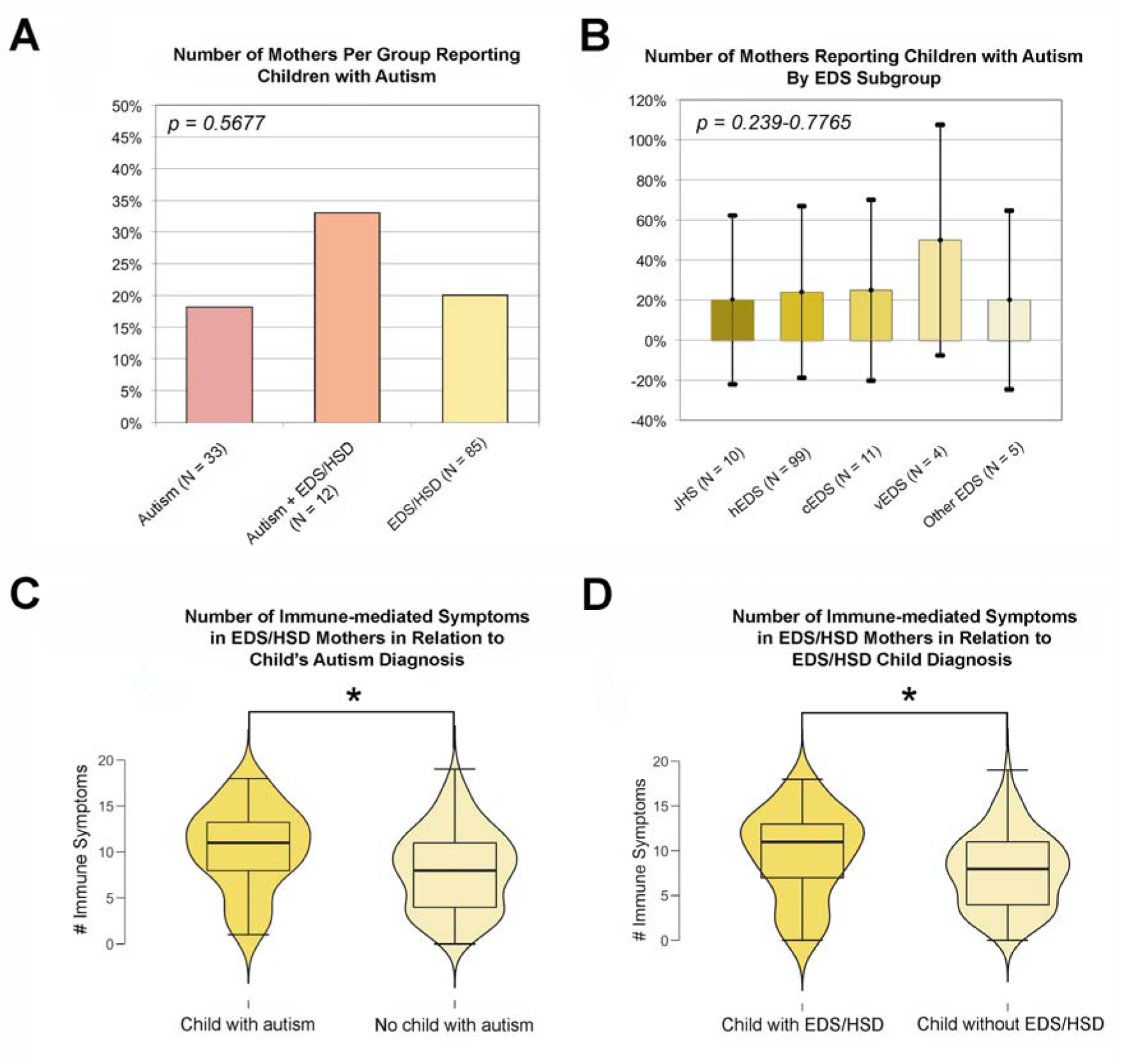
Autism and EDS/HSD in the Offspring. (A) Reported rates of autism diagnosis in children of mothers with clinical syndromes of interest. (B) Reported rates of EDS/HSD diagnosis in children of mothers with clinical syndromes of interest. (C) Rates of immune-mediated symptoms in mothers with EDS/HSD in relation to the child’s autism diagnosis. (D) Rates of immune-mediated symptoms in mothers with EDS/HSD in relation to the child’s EDS/HSD diagnosis.

More intriguingly, the distribution of immune-mediated symptoms in EDS/HSD mothers who had autistic children significantly differed from those with non-autistic children, with the former reporting on average 28% more symptoms than the latter [mean = 10.4 symptoms (Sx) > 8.1 Sx, *t* = −2.51, *p* = 0.015] (Fig. 2c). Surprisingly, the distribution of immune symptoms between EDS/HSD mothers who had EDS/HSD children and those without also significantly differed, with the former reporting about 25% more symptoms than the latter (mean = 9.6 Sx > 7.7 Sx) [*W* = 1496, *p* = 0.0145] (Fig. 2d). However, neither of these groups experienced significantly increased autonomic or endocrine-mediated symptoms [*W* = 1321-1760, *p* = 0.0945-0.2356] (1) (see Supplementary File 2, Fig. 1). Therefore, the etiologies of both EDS/HSD and autism seem to share links with the maternal immune system in particular (Garay et al., 2013; McDougle, 2018). (For more results involving immune dysregulation, see Supplementary File 2, Fig. 2-3.)

### Other Ehlers-Danlos-related Symptomology

We collected data on additional symptomology associated with EDS/HSD, in order to compare relative frequencies across groups (Table 2). (Due to small numbers within the male EDS/HSD subgroup, all results reported here concern females only unless otherwise stated.) In terms of hypermobility, joint dislocations/subluxations, skin fragility, chronic pain, general clumsiness, and chronic fatigue, as expected the EDS/HSD group reported significantly more symptoms compared to both the autistic and control groups [all *p* > 0.001-0.0022] (Fig. 3). The autism group likewise differed from controls in terms of clumsiness, chronic pain, chronic fatigue, and skin fragility [all *p* ≤ 0.002], but did not differ as to hypermobility and dislocations/subluxations [all *p* = 0.136-0.6568]. Males with autism also significantly differed from control males in the same regards [all *p* = 0.028-0.042]. These data suggest that some forms of chronic pain, tissue fragility, and fatigue may be of concern even in autism unassociated with EDS/HSD.

**Figure 3.**
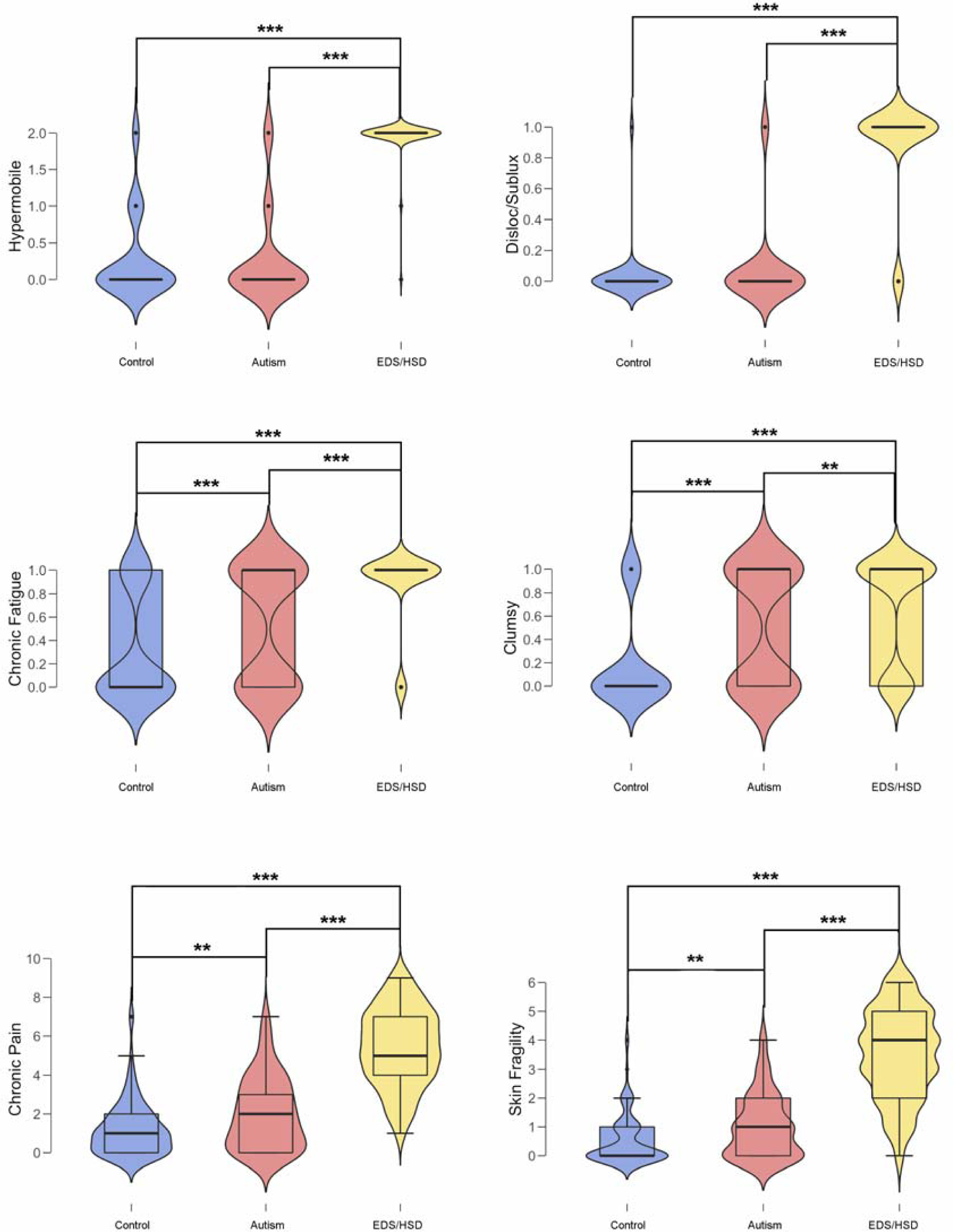
EDS/HSD-associated Symptoms. Reported rates of hypermobility, dislocations/subluxations, chronic fatigue, clumsiness, chronic pain, and skin fragility according to group.

As expected, we also found that the distribution of vascular complications, such as strokes, aneurysms, and hemorrhages, significantly differed across groups and were far more common in the EDS/HSD group (21%), while they did not differ between autism and controls (2% vs. 4%) [*X*^*2*^ = 24.809, *p* < 0.001]. Likewise, the log odds of head injuries and proportions of Chiari malformations differed significantly across groups, and were unusually high in the EDS/HSD group, findings which have previously been reported in the EDS literature [head injury: *X*^*2*^ = 19.086, *p* < 0.0001; Chiari: Fisher’s Exact Test *p* = 0.002] (Hamonet et al., 2016; Malfait et al., 2010; Castori et al., 2010) (Table 2).

### Neurodevelopmental and Psychiatric Conditions

Both autism and EDS/HSD are sometimes associated with other neurodevelopmental and/or psychiatric conditions. When asked to report on comorbid diagnoses of attention deficit/hyperactivity disorder (ADHD), anxiety disorders, mood disorders, and post traumatic stress disorder (PTSD), we found that reports significantly differed by group, although this was largely driven by clinical groups versus controls [*X*^*2*^ = 10.6538-16.402, *p* < 0.0066-0.0138]. In fact, when analyzed individually, EDS/HSD and autism did not significantly differ from one another, although individuals with autism did report more mood disorders than those with EDS/HSD (47% vs. 29%). This suggests that neurodevelopmental and psychiatric comorbidities are similar across our two clinical groups and that some features of EDS/HSD may be neurodevelopmental in origin. We also found that log odds of comorbidity did not differ significantly across the EDS/HSD subgroups, indicating that they are all similarly affected and that neurodevelopmental and psychiatric conditions do not cluster within select forms of EDS [*X*^*2*^ = 1.0026-3.017, all adjusted *p* = 0.8006] (Bulbena-Cabré & Bulbena, 2018).

## DISCUSSION

This study attempted to investigate a variety of health issues across EDS/HSD, autism, and controls. Much of the survey design was guided by previous EDS/HSD research (Casanova et al., 2018). Given the apparent (though still poorly delineated) comorbidity between EDS/HSD and autism, we felt it was important to determine to what extent EDS/HSD issues affect the autism spectrum.

In summarizing the data presented here, those with autism (without EDS/HSD) appear to have significant issues concerning immune/autonomic/endocrine dysregulation, connective tissue (skin) durability, chronic pain, and fatigue. In addition, autistic women experience more endocrine disorders than sex-matched controls. All of these symptoms appear to be less extreme than those seen in individuals with EDS/HSD, which may share striking comorbidity with autism. In contrast, issues such as hypermobility, dislocations/subluxations, vascular complications, chiari malformation, and head injury all appear to be largely specific to the EDS/HSD diagnosis.

Autism seems to occur alongside EDS/HSD with some frequency, although we cannot estimate comorbidity rates in this study due to potential recruitment bias. There is also evidence to suggest that maternal EDS/HSD could be a risk factor for the development of autism and EDS/HSD in the offspring, although, once again, more research is needed to address this possibility. In particular, our data indicate that the maternal immune system may play a regulatory role, a finding that echoes the general autism literature (Patterson, 2011).

## Limitations

As with all survey studies, the veracity and applicability of our data is a reflection of the reliability of reporting of our respondents. Because our findings generally agree with those previously reported throughout the EDS/HSD and autism literature, this lends support to the data and the conclusions presented here, although clearly more clinical research is needed (Seneviratne et al., 2017; Kushki et al., 2013; De Wandele et al., 2014; Careaga et al., 2017; Ingudomnukul et al., 2007).

Recent changes in hEDS nosology also make it challenging to accurately assign participants to the EDS versus HSD subheadings as many will have received diagnoses prior to the latest diagnostic update. For this reason all respondents reporting a form of EDS, G-HSD, or JHS have been subsumed under the general “EDS/HSD” heading for most analyses. Given evidence of their relatedness both in this and previous studies, as well as the musculoskeletal impairment reported by our sample of respondents, we feel such a compilation is justified.

Another limitation of some of our EDS/HSD data involves small sample sizes. While the number of hEDS participants was substantial, the rarer forms of EDS, such as vEDS, were especially small in number and it is therefore difficult to draw conclusions about their responses, except to say that with current numbers the vEDS data do not appear to differ substantially from the other EDS/HSD subgroups. While this must be interpreted cautiously, it does suggest that people with vEDS experience many of the same secondary clinical phenotypes as the more common forms of EDS/HSD, reinforcing concepts of a common Ehlers-Danlos syndrome despite genetic heterogeneity. Further research will continue to test this possibility.

Finally, as has been mentioned previously, we cannot address comorbidity and inheritance rates between EDS/HSD and autism due to the ways in which participants were recruited. It is therefore possible and perhaps likely that individuals with comorbid EDS/HSD and autism diagnoses or EDS/HSD parents with autistic children were more drawn to participate. For these reasons we have simply described our study population qualitatively.

## CONCLUSIONS

Although we cannot estimate comorbidity rates, the data presented here suggest at least a small portion of the autism spectrum may fulfill criteria for EDS/HSD. Prevalence rates of EDS are poorly researched, with current data estimating the condition affects 1:5,000 individuals. However, current clinical expertise suggests this is an underestimation and that it is far more common (EDS Society, 2018). In summary, these data suggest an intriguing possibility that some forms of autism are HCTDs. Our laboratory continues to work to address this possibility and to help improve the lives of those with these spectrum conditions dealing with chronic illness.

## Supporting information

Supplementary File - Survey

Supplementary File - Statistics & Additional Results

Supplementary File - Study Data

## ACKNOWLEDGMENTS

We would like to thank the Ehlers-Danlos syndrome/hypermobility spectrum disorder and autistic online communities who took part in and actively supported this research. We would also like to thank the online websites that allowed us to advertise this study.

## COMPLIANCE WITH ETHICAL STANDARDS

All procedures performed in studies involving human participants were in accordance with the ethical standards of the Greenville Health System’s Institutional Review Board (ID: Pro00073030) and with the 1964 Helsinki declaration and its later amendments or comparable ethical standards. All participants completed a waiver of consent in order to participate in this online study.

## AVAILABILITY OF DATA AND MATERIALS

Full raw data, as well as select statistical and descriptive results, produced in this study are available within the Supplementary Files provided with the manuscript. For any data or statistical results not available therein, the primary author (ELC) can be contacted for full access.

