## Supplementary File - Survey for "Immune, Autonomic, and Endocrine Dysregulation in Autism and Ehlers-Danlos Syndrome/Hypermobility Spectrum Disorders Versus Unaffected Controls"

### Expanding Our Understanding of Symptoms Associated with Autism Spectrum Disorder & Ehlers-Danlos Syndrome

**Page exit logic:** Skip / Disqualify Logic
**IF:** #1 Question "Do you agree to participate in this survey study? Please read the options below carefully." is one of the following answers ("No, I do not consent.") **THEN:** Jump to page 13 - Thank You! Flag response as complete

ID 5

RESEARCH STUDY INFORMATION

#### Study to be Conducted at:

Greenville Health System
Patewood Medical Campus
200 Patewood Dr.
Suite 200A
Greenville, SC 29615

#### Principal Investigator:

Manuel Casanova, MD

#### Co-investigator:

Emily Casanova, PhD

#### Introduction:

You are being asked to participate in a research study. The Institutional Review Board of the Greenville Health System has reviewed this study for the protection of the rights of human participants in research studies, in accordance with federal and state regulations. However, before you choose to be a research participant, it is important that you read the following information and ask as many questions as necessary to be sure that you understand what your participation will involve.

#### Purpose and Procedures:

We are looking for adults 25 years of age or older (or legal guardians of adults who fulfill those same criteria) to participate in a survey study. The purpose of this research is to expand our understanding of the types and frequency of certain medical symptoms experienced in Autism Spectrum Disorder (ASD), Ehlers-Danlos Syndrome (EDS)/Joint Hypermobility

Syndrome (JHS), and their families. We are also interested in studying the same medical issues in the general population. **You are being asked to participate because you are one of the following:**

1. *An adult 25 years of age or older with a diagnosis of ASD and/or EDS/JHS.*
2. *A legal guardian of an adult 25 years of age or older who would like to answer the survey on his/her behalf.*
3. *An adult 25 years of age or older who has a close family member with ASD and/or EDS/JHS.*
4. *An adult 25 years of age or older who would like to participate as a control case for this study.*

We plan to enroll at least several hundred people in this study. Your participation will involve answering an online survey to the best of your abilities. This should take approximately 15-25 minutes.

#### Possible Risks and Benefits:

There are no known medical risks related to participation in this study. The greatest risk is the possible release of your personal information with the investigators. However, this survey is anonymous and we will not be collecting any identifying information. Your survey answers are considered confidential, but absolute confidentiality cannot be guaranteed. This study may result in presentations and publications. There are no direct benefits to you that would result from your participation in the study.

Although you will not receive compensation for participating in this study, this research may help us to understand ASD and EDS better. This knowledge can help scientists and doctors develop better treatments for the varied symptoms associated with these conditions that may affect you and/or people you know.

#### Voluntary Participation:

Participation in this study is completely voluntary (your choice). You may refuse to participate or withdraw prior to submitting your survey. If you refuse to participate or stop the survey, you will not be penalized or lose any benefits. However, because your submission is anonymous, we will not be able to remove your data from the study afterwards. So please be certain you want to participate prior to submission. Your decision will not affect your relationship with the investigators or any relationship you may have with the Greenville Health System.

#### Contact for Questions:

For more information concerning this study and research-related risks or injuries, or to give comments or express concerns or complaints, you may contact the principal investigator, *Manuel Casanova*, at (864) 454-4595. You may also contact a representative of the Institutional Review Board of the Greenville Health System for information regarding your rights as a participant involved in a research study or to give comments or express concerns, complaints, or offer input. You may obtain the name and number of this person by calling (864) 455-8997.

An additional survey about your experience with this informed consent process is located at the following website: https://www.surveymonkey.com/s/T5C86P8 Participation in the above survey is completely anonymous and voluntary and will not affect your relationship with the Greenville Health System. If you would like to have a paper copy of this survey, please tell the principle investigator.

ID 6

1. Do you agree to participate in this survey study? Please read the options below carefully. *

- Yes, I am an adult 25 years of age or older and would like to participate in this study.
- Yes, I am the legal guardian of an adult 25 years of age or older and would like to participate in this study on his/her behalf.
- No, I do not consent.

**Background Information**

LOGIC Show/hide trigger exists.

ID 172

2. Are you participating on your own behalf or as a legal guardian on behalf of your **adult** ward? *

- Self
- Legal guardian

LOGIC Hidden unless: #2 Question "Are you participating on your own behalf or as a legal guardian on behalf of your **adult** ward?" is one of the following answers ("Legal guardian")

ID 173

**An important note:** Please answer the following questions on behalf of your ward. Whenever a question refers to the reader as "you," this refers to your ward. Some questions, such as parent medical history, may concern you directly. Please answer all questions on behalf of your ward to the best of your abilities.

LOGIC Show/hide trigger exists.

ID 8

3. Sex: *

- ○ Male
- ○ Female
- ○ Other, please specify: ________ *

ID 9

4. Current age in years. (Please do NOT provide birth date.) *

________

ID 118

5. In which country were you born? *

________

ID 145

6. If you live in the United States of America and would be willing to share information, in which state do you live? (This question is optional. We're hoping to determine whether certain symptoms are more common or severe in specific regions of the country.)

ID 50

7. What is your ancestry? Below are groups of people who have related ancestries. Please check all that you think apply to you. If you're not certain, give best estimates or select "Don't know." *

- ☐ Sub-saharan African (West, East, Central, and South Africa)
- ☐ Middle Eastern & Northern African
- ☐ East Asian & Native American
- ☐ South Asian (India, Pakistan, Afghanistan)
- ☐ Northwestern European (British Isles, Germany, France, Scandinavia)
- ☐ Eastern European (Poland, Russia, etc.)
- ☐ Southern European (Spain, Portugal, Italy, Greece)
- ☐ Don't know

ID 51

8. Do you have anything else you'd like to clarify regarding your ancestry?

ID 141

9. Are you adopted? *

- Yes
- No
- I prefer not to answer

LOGIC Show/hide trigger exists.

ID 174

10. Were you born premature? *

- Yes
- No
- Don't know

LOGIC Hidden unless: #10 Question "Were you born premature?" is one of the following answers ("Yes")

ID 144

11. How premature were you? *

- Moderate to late preterm (32 to <37 weeks gestational age)
- Very preterm (28 to <32 weeks gestational age)
- Extremely preterm (<28 weeks gestational age)
- Not sure

LOGIC Show/hide trigger exists.

ID 138

12. Do you have a professional diagnosis of Autism Spectrum Disorder (ASD) or a related diagnosis (e.g., Asperger's Syndrome)? If so, please write the diagnosis in the box provided. *

- Yes: ______ *
- Suspected
- No
- Other, please explain: ______ *

LOGIC Hidden unless: #12 Question "Do you have a professional diagnosis of Autism Spectrum Disorder (ASD) or a related diagnosis (e.g., Asperger's Syndrome)? If so, please write the diagnosis in the box provided." is one of the following answers ("Yes:")

ID 193

13. Can you describe where, when, and by whom you were diagnosed with ASD? If you don't recall all the information, please describe to the best of your abilities. *

LOGIC Show/hide trigger exists.

ID 139

14. Do any of your 1st degree **biological** relatives (parent, sibling, child) have an ASD diagnosis? *

- Yes
- Suspected
- No
- Don't know
- Other, please explain: *

LOGIC Hidden unless: #14 Question "Do any of your 1st degree **biological** relatives (parent, sibling, child) have an ASD diagnosis?" is one of the following answers ("Yes")

ID 189

15. Which of your 1st degree biological relatives is **diagnosed** with an ASD? Please check all that apply. *

- Father
- Mother
- Brother
- Sister
- Son
- Daughter

LOGIC Hidden unless: #14 Question "Do any of your 1st degree **biological** relatives (parent, sibling, child) have an ASD diagnosis?" is one of the following answers ("Yes")

ID 200

16. Please describe to the best of your abilities where, when, and by whom your 1st degree biological relative was diagnosed with an ASD.

LOGIC Show/hide trigger exists.

ID 10

17. Do you have a professional diagnosis of Ehlers-Danlos Syndrome (EDS) or Joint Hypermobility Syndrome (JHS)? *

- Yes
- Suspected
- No
- Other, please explain: *

LOGIC Show/hide trigger exists. Hidden unless: #17 Question "Do you have a professional diagnosis of Ehlers-Danlos Syndrome (EDS) or Joint Hypermobility Syndrome (JHS)?" is one of the following answers ("Yes")

ID 12

18. What type of EDS are you diagnosed with? If you are not diagnosed with EDS but with Joint Hypermobility Syndrome (JHS), please click the option below. *

- EDS, Hypermobility Type
- Joint Hypermobility Syndrome (JHS)
- EDS, Classic Type
- EDS, Vascular Type
- EDS, Kyphoscoliosis Type
- EDS, Arthrocalasia Type
- EDS, Dermatosparaxis Type
- EDS, Tenascin-X Deficiency Type
- Other, please specify: *

LOGIC Hidden unless: #17 Question "Do you have a professional diagnosis of Ehlers-Danlos Syndrome (EDS) or Joint Hypermobility Syndrome (JHS)?" is one of the following answers ("Yes")

ID 194

19. Can you describe where, when, and by whom you were diagnosed with EDS/JHS? If you don't recall all the information, please describe to the best of your abilities. *

LOGIC Hidden unless: #17 Question "Do you have a professional diagnosis of Ehlers-Danlos Syndrome (EDS) or Joint Hypermobility Syndrome (JHS)?" is one of the following answers ("Suspected")

ID 13

20. What type of EDS do you suspect or been told you may have? *

- EDS, Hypermobility Type
- Joint Hypermobility Syndrome (JHS)
- EDS, Classic Type
- EDS, Vascular Type
- EDS, Kyphoscoliosis Type
- EDS, Arthrocalasia Type
- EDS, Dermatosparaxis Type
- EDS, Tenascin-X Deficient Type
- Other, please specify: *

LOGIC Show/hide trigger exists. Hidden unless: #18 Question "What type of EDS are you diagnosed with? If you are not diagnosed with EDS but with Joint Hypermobility Syndrome (JHS), please click the option below." is one of the following answers ("EDS, Hypermobility Type","Joint Hypermobility Syndrome (JHS)","EDS, Classic Type","EDS, Vascular Type","EDS, Kyphoscoliosis Type","EDS, Arthrocalasia Type","EDS, Dermatosparaxis Type","EDS, Tenascin-X Deficiency Type","Other, please specify:")

ID 22

21. Do you have a genetic mutation(s) that a geneticist has identified as potentially/probably responsible for your EDS/JHS? *

- Yes
- No
- Don't know
- Other, please explain: *

LOGIC Hidden unless: #21 Question "Do you have a genetic mutation(s) that a geneticist has identified as potentially/probably responsible for your EDS/JHS?" is one of the following answers ("Yes")

ID 23

22. If you know the mutation please list it here. If you don't know/recall, please say so. *

LOGIC Show/hide trigger exists.
ID 202
23. Do any of your 1st degree **biological** relatives (parent, sibling, child) have a diagnosis of EDS or JHS? Please check all that apply. *

- ☐ EDS
- ☐ JHS
- ☐ Suspected EDS/JHS
- ☐ No
- ☐ Don't know
- ☐ Other, please explain:

LOGIC Hidden unless: #23 Question "Do any of your 1st degree **biological** relatives (parent, sibling, child) have a diagnosis of EDS or JHS? Please check all that apply." is one of the following answers ("EDS","JHS")
ID 191
24. Which of your 1st degree biological relatives has a **diagnosis** of EDS/JHS? Please check all that apply. *

- ☐ Father
- ☐ Mother
- ☐ Brother
- ☐ Sister
- ☐ Son
- ☐ Daughter

LOGIC Hidden unless: #23 Question "Do any of your 1st degree **biological** relatives (parent, sibling, child) have a diagnosis of EDS or JHS? Please check all that apply." is one of the following answers ("EDS","JHS")

ID 201

25. Please describe to the best of your abilities where, when, and by whom your 1st degree biological relative was diagnosed with EDS/JHS. Also, please specify his/her diagnosis.

ID 161

26. Do you have a diagnosis of any kind of heritable or teratogenic (environmentally-induced) syndrome? For example, Fragile X Syndrome is a genetic condition due to mutations in the *FMR1* gene. Meanwhile, Fetal Alcohol Syndrome is an environmentally-induced condition due to maternal alcohol consumption during pregnancy. Please indicate the syndrome(s) in the box provided. *

- Yes: *
- No
- Don't know
- Other, please explain: *

#### The Brain

ID 175

27. Please estimate your intelligence quotient (IQ). If you've never had an IQ test or don't remember the results, give your best guess. *

- Low (<85)
- Average (86-115)
- High (116+)

ID 41

28. Do you have a professional diagnosis of another neurodevelopmental, psychiatric, or neurological condition (e.g., Generalized Anxiety Disorder, ADHD, etc) not covered in the previous questions? If so, please list the diagnoses. *

- Yes: ______ *
- Suspected: ______ *
- No
- Don't know
- Other, please specify: ______ *

LOGIC Show/hide trigger exists.

ID 49

29. Do you have a history of epilepsy (either current or in the past)? *

- Yes, epilepsy with no known cause
- Yes, epilepsy following head trauma, infection, or another identifiable cause
- No
- Don't know
- Other, please explain: *

LOGIC Hidden unless: #29 Question "Do you have a history of epilepsy (either current or in the past)?" is one of the following answers ("Yes, epilepsy with no known cause","Yes, epilepsy following head trauma, infection, or another identifiable cause")

ID 203

30. At roughly what age did you develop epilepsy?

ID 187

31. Have any of your 1st degree **biological** relatives (parent, sibling, child) had epilepsy? *

- Yes
- No
- Don't know
- Other, please explain: *

ID 155

32. As an infant or young child, did you ever experience febrile (fever-related) seizures? *

- Yes
- No
- Don't know
- Other, please explain: ____________ *

LOGIC Hidden unless: (#32 Question "As an infant or young child, did you ever experience febrile (fever-related) seizures?" is one of the following answers ("Yes") AND #29 Question "Do you have a history of epilepsy (either current or in the past)?" is one of the following answers ("Yes, epilepsy with no known cause","Yes, epilepsy following head trauma, infection, or another identifiable cause"))

ID 156

33. Did these febrile seizures transition into epilepsy or did your epilepsy seem to arise independently? *

- The febrile seizures transitioned into epilepsy
- My epilepsy arose independent of the febrile seizures
- Not sure
- Other, please explain: ____________ *

ID 131

34. Have you ever experienced a traumatic brain injury (TBI) in relation to an acute physical trauma, such as a car accident, an aneurysm, or a stroke? If so, please describe the injury in the space provided. *

- Yes: *
- No
- Don't know/unsure
- Other, please explain: *

ID 128

35. Do you now or have you ever experienced migraines? *

- Yes
- No
- Don't know
- Other, please explain: *

ID 129

36. Do you experience other chronic problems with non-migraine headaches? *

- Yes
- Sometimes
- No
- Don't know
- Other, please explain: *

ID 90

37. As a rough estimate of general head size, when you wear hats what size do you normally wear (relative to your sex)? *

- Extra-large
- Large
- Medium
- Small
- Don't know

#### Skin & Joints

LOGIC Show/hide trigger exists.

ID 14

38. Are you hypermobile (double-jointed)? *

- Yes, across two or more types of joints (e.g., wrists, knees, elbows, etc.).
- Yes, in a single type of joint (e.g., elbows).
- No
- Don't know
- Other, please explain: *

LOGIC Hidden unless: #38 Question "Are you hypermobile (double-jointed)?" is one of the following answers ("Yes, across two or more types of joints (e.g., wrists, knees, elbows, etc.).","Yes, in a single type of joint (e.g., elbows).")

ID 85

39. In which joints are your hypermobile? *

LOGIC Hidden unless: #17 Question "Do you have a professional diagnosis of Ehlers-Danlos Syndrome (EDS) or Joint Hypermobility Syndrome (JHS)?" is one of the following answers ("Yes")

ID 209

40. Have you been told by a doctor you have a "Marfanoid habitus"? *

- Yes
- Maybe
- No
- Not sure
- Other, please explain: *

LOGIC Show/hide trigger exists. Hidden unless: #17 Question "Do you have a professional diagnosis of Ehlers-Danlos Syndrome (EDS) or Joint Hypermobility Syndrome (JHS)?" is one of the following answers ("Yes")

ID 205

41. Do you use any mobility devices such as a wheelchair, walker, or cane because of the effects of your EDS/JHS? If so, please list the device(s) you use:

- Always: *
- Sometimes: *
- Rarely
- Never

LOGIC Hidden unless: #41 Question "Do you use any mobility devices such as a wheelchair, walker, or cane because of the effects of your EDS/JHS? If so, please list the device(s) you use:" is one of the following answers ("Always:","Sometimes:")

ID 207

42. Did you start using any of these mobility devices prior to the age of 50?

- ○ Yes
- ○ No
- ○ Don't know
- ○ Other, please explain: ______ *

ID 176

43. Do you consider yourself clumsy or uncoordinated? *

- ○ Yes
- ○ Sometimes
- ○ No
- ○ Don't know
- ○ Other, please explain: ______ *

LOGIC Show/hide trigger exists.

ID 15

44. Please indicate which, if any, of these types of chronic pain you experience that is unusual for your age (or at one time was) and isn't the result of an acute injury (e.g., a car accident). Please indicate to the right the approximate age around which you started experiencing this pain. *

- [ ] Joint pain (arthralgia): ______ *
- [ ] Muscle pain (myalgia): ______ *
- [ ] Arthritis: ______ *
- [ ] Fibryomyalgia: ______ *
- [ ] Temporomandibular Joint (TMJ) pain: ______ *
- [ ] Neuropathy-induced pain: ______ *
- [ ] Pain due to endometriosis: ______ *
- [ ] Pain due to another organ (e.g., bladder, prostate, etc.): ______ *
- [ ] Other, please describe: ______ *
- [ ] None of the above

LOGIC Hidden unless: #44 Question "Please indicate which, if any, of these types of chronic pain you experience that is unusual for your age (or at one time was) and isn't the result of an acute injury (e.g., a car accident). Please indicate to the right the approximate age around which you started experiencing this pain." is one of the following answers ("Joint pain (arthralgia):","Muscle pain (myalgia):","Arthritis:","Fibryomyalgia:","Temporomandibular Joint (TMJ) pain:","Neuropathy-induced pain:","Pain due to endometriosis:","Pain due to another organ (e.g., bladder, prostate, etc.):","Other, please describe:")

ID 163

45. Did you ever experience a stressful event (e.g., emotional or physical) after which your pain unexpectedly worsened? For example, you may have experienced flares of TMJ following surgery for an unrelated issue. If so, please describe what happened. If not, please say so. *

ID 146

46. Do you have a diagnosis of Chronic Fatigue Syndrome? *

- Yes
- Suspected
- No
- Don't know
- Other, please explain: *

ID 71

47. Do you experience recurrent dislocations or subluxations (partial dislocation) of joints? *

- Yes
- No
- Don't know
- Other, please explain: *

ID 135

48. Were you born with any dislocations (e.g., hips)? If so, please describe the dislocations in the space provided. *

- Yes: *
- No
- Don't know
- Other, please explain: *

ID 121

49. Do you have foot problems, such as recurrent plantar fasciitis, flat feet, high arches, pronation (rolling ankles), or poorly healing foot injuries? *

- Yes
- No
- Don't know
- Other, please explain:

*

ID 16

50. Do you have any chronic pain not addressed by the previous questions?

ID 17

51. Do you have any of the following issues? Please check all that apply. If you have none of these, please check, "None of the above." *

- Bruise easily
- Have bleeds that are difficult to control
- Scar easily
- "Cigarette paper" scars
- Unusually elastic skin
- Slow wound healing
- None of the above

ID 132

52. Have you been told throughout much of your life that you look younger than your age? *

- Often
- Sometimes
- Rarely
- Don't know
- Other, please explain:

  *

ID 123

53. Have you been told by a health professional that you have extra bones, such as an extra rib or an accessory navicular bone? If so, please list the bones or if you don't know their names describe them as best you can. *

- Yes:

  *

- No
- Don't know
- Other, please explain:

  *

ID 124

54. Have you been told by a health professional that you have congenitally missing bones? If so, please list the bones or if you don't know their names describe them as best you can. *

- Yes: ______ *
- No
- Don't know
- Other, please explain: ______ *

ID 87

55. Do you have or did you have at one time dental crowding/crooked teeth? *

- Yes
- No
- Don't know
- Other, please explain: ______ *

ID 91

56. Have you ever had a slipped spinal disc? *

- Yes
- Suspected
- No
- Don't know
- Other, please explain:

ID 134

57. Do you have any of the following? If not, please select "None of the above." *

- Scoliosis (sideways curvature of the spine)
- Kyphosis (hunchback)
- Scleral fragility (rupture of the globe of the eye)
- Respiratory problems due to kyphosciolosis
- None of the above

ID 89

58. Have you had any arterial, intestinal, or uterine ruptures not otherwise due to blunt trauma (e.g., surgery)? *

- Yes
- No
- Don't know
- Other, please explain:

*

ID 84

59. Do you have a chiari malformation? *

- Yes
- Suspected
- No
- Don't know
- Other, please explain: *

ID 122

60. Do you have any types of polyps, cysts, or fibroids? If so, what type(s)? *

- Yes: *
- No
- Don't know
- Other, please specify: *

LOGIC Hidden unless: #44 Question "Please indicate which, if any, of these types of chronic pain you experience that is unusual for your age (or at one time was) and isn't the result of an acute injury (e.g., a car accident). Please indicate to the right the approximate age around which you started experiencing this pain." is one of the following answers ("Joint pain (arthralgia):","Muscle pain (myalgia):","Arthritis:","Fibryomyalgia:","Temporomandibular Joint (TMJ) pain:","Neuropathy-induced pain:","Pain due to endometriosis:","Pain due to another organ (e.g., bladder, prostate, etc.):","Other, please describe:")

ID 167

61. On a typical day, how would you rate your general level of pain? *

| No pain | Mild Pain | Moderate Pain | Severe Pain | Very Severe Pain | Worst Pain Possible |
|---|---|---|---|---|---|
| ○ | ○ | ○ | ○ | ○ | ○ |

#### Cardiovascular

ID 86

62. Do you have mitral valve prolapse *

- ○ Yes
- ○ No
- ○ Don't know
- ○ Other, please explain: *

ID 133

63. Have you ever been diagnosed with any of the following? Please check all that apply. If you have none, please check, "None of the above." *

- ☐ Spontaneous arterial dissections (tears within arteries)
- ☐ Cavernous sinus fistula (abnormal connection between the carotid artery in the neck and veins at the back of the eye)
- ☐ Aneurysm
- ☐ Brain hemorrhage
- ☐ Stroke
- ☐ Other vascular complication not listed
- ☐ None of the above

#### Eyes

ID 116

64. Do you have (or have you ever had) any of the following? Please check all that apply. If you have none of these, please click, "None of the above." *

- ☐ Myopia (short-sightedness)
- ☐ Frequent, easily agitated eye strain
- ☐ Retinal detachment
- ☐ Ptosis (droopy eyelid)
- ☐ Blue or gray sclera (coloration within the whites of the eyes)
- ☐ Astigmatism
- ☐ Keratoconus (bulging, cone-shaped cornea due to corneal thinning)
- ☐ Chronic dry eye
- ☐ Strabismus (improper alignment of the eyes)
- ☐ None of the above

ID 142

65. How much time to you usually spend on the computer per day? *

- Less than 1 hour
- 1-2 hours
- 2-3 hours
- 4+ hours

#### Postural Orthostatic Tachycardia Syndrome (POTS)

ID 48

66. Do you have a diagnosis of Postural Orthostatic Tachycardia Syndrome (POTS)? *

- Yes
- Suspected, but not officially diagnosed
- No
- Don't know
- Other, please explain: *

ID 44

67. Do you experience any of the following on a frequent and regular basis? Please check all that apply. If you have none of these, please click, "None of the above." *

- ☐ Chronic fatigue or transient unexplained tiredness
- ☐ Dizziness, light-headedness, vertigo, especially upon standing
- ☐ Fainting
- ☐ Brain fog
- ☐ Heart palpitations
- ☐ Chest pains
- ☐ Shortness of breath, difficulty taking a deep breath
- ☐ Weakness
- ☐ Headaches (non-migraine)
- ☐ Abnormal sweating (too much or too little)
- ☐ Gastrointestinal distress (abdominal pain, nausea, vomiting, diarrhea, constipation)
- ☐ Muscle tremors
- ☐ Bladder dysfunction/incontinence
- ☐ Insomnia
- ☐ Free-floating (unexplained) anxiety
- ☐ None of the above

ID 164

68. Does your blood pressure fluctuate unusually? For instance, does it show up high in the physician's office but when tested at home it's normal or even low? *

- Yes
- Sometimes
- No
- Don't know
- Other, please explain:

*

ID 158

69. Do you regularly run a low or high body temperature or feel like you often have trouble regulating your temperature? For instance, do you frequently feel too cold or too hot, or even vacillate between extremes at different times of the day? (These changes should not be due to conditions such as menopause.) *

- Yes
- Maybe
- No
- Don't know
- Other, please explain:

*

ID 143

70. Have you ever had a heart attack or been diagnosed with heart disease? *

- Yes
- No
- Don't know
- Other, please explain:

*

#### The Immune System

ID 157

71. Do you have a diagnosis of either Mast Cell Activation Syndrome (MCAS) or Systemic Mastocytosis? *

- Diagnosed with MCAS
- Suspect MCAS
- Diagnosed with systemic mastocytosis
- Suspect systemic mastocytosis
- None of the above
- Other, please explain:

*

ID 24

72. Any history of hospitalization due to **infection** during the childhood, teenage years, or adulthood? Check all that apply. *

- ☐ Childhood (0-9 yrs)
- ☐ Teen (10-17)
- ☐ Adult (18 yrs & older)
- ☐ No hospitalizations due to infection

LOGIC Show/hide trigger exists.

ID 28

73. Any chronic history of the following conditions? Please indicate your answers in the checkboxes below. If you have none of these, please check, "None of the above" at the end of the list. *

- ☐ Anaphylaxis
- ☐ Respiratory allergies
- ☐ Asthma
- ☐ Frequent ear infections in childhood
- ☐ Frequent ear infections in adulthood
- ☐ Frequent rhinitis (inflammation of the nose)
- ☐ Frequent sinusitis (inflammation of the sinuses)
- ☐ Dry, itchy eyes
- ☐ Skin allergy
- ☐ Eczema (dry, inflamed skin)
- ☐ Hives/rash
- ☐ Food allergy
- ☐ Severe or unusual physical reactions to medications (anaphylaxis, severe drowsiness, rash, etc.)
- ☐ Severe or unusual physical reactions to environmental chemicals (runny nose, coughing, asthma, rash, etc.)
- ☐ Severe or unusual physical reactions to emotional or physical stress
- ☐ Frequent urinary tracts infections or inflammation
- ☐ Frequent vaginal infections or inflammation
- ☐ Stomach ulcers
- ☐ Reflux
- ☐ Transiently swollen lymph nodes (not due to infection or cancer)
- ☐ None of the above

LOGIC Hidden unless: #73 Question "Any chronic history of the following conditions? Please indicate your answers in the checkboxes below. If you have none of these, please check, "None of the above" at the end of the list." is one of the following answers ("Food allergy")

ID 154

74. What food(s) are/were you allergic to?

LOGIC Hidden unless: #73 Question "Any chronic history of the following conditions? Please indicate your answers in the checkboxes below. If you have none of these, please check, "None of the above" at the end of the list." is one of the following answers ("Respiratory allergies","Skin allergy","Food allergy")

ID 97

75. Please describe any allergies you may have and your daily and/or seasonal experience with those allergies. *

ID 160

76. Have you had any diagnoses of chronic inflammation of a specific organ, such as "pancreatitis" or "interstitial cystitis"? Often these conditions end with the suffix *-itis.* Please indicate in the box what diagnoses you've received. *

- Yes: *
- No
- Don't know
- Other, please explain: *

ID 38

77. Are you diagnosed with an autoimmune disease, such as Diabetes I or Rheumatoid Arthritis? If so, what is it? *

- Yes: *
- Suspected: *
- No
- Don't know

LOGIC Show/hide trigger exists.

ID 47

78. Have any of your **biological** 1st degree relatives (parent, sibling, child) been diagnosed with an autoimmune condition? *

- Yes
- Suspect
- No
- Don't know
- Other, please explain: *

LOGIC Hidden unless: #78 Question "Have any of your **biological** 1st degree relatives (parent, sibling, child) been diagnosed with an autoimmune condition?" is one of the following answers ("Yes")

ID 185

79. Please check which family members have/had an autoimmune disease and list the type. If you're not certain, please answer to the best of your abilities. *

- [ ] Father: ______ *
- [ ] Mother: ______ *
- [ ] Brother: ______ *
- [ ] Sister: ______ *
- [ ] Son: ______ *
- [ ] Daughter: ______ *

ID 165

80. Have you ever gotten sick closely following a vaccination? If so and you recall the type of vaccine, please write it in the space provided. *

- Yes: *
- No
- Don't know
- Other, please explain: *

ID 166

81. Have you ever had problems developing immunity? For instance, did you require re-vaccination after the normal number of rounds or did you experience an illness like chickenpox more than once? Please describe in the box provided, *

- Yes: *
- No
- Don't know
- Other, please explain: *

LOGIC Show/hide trigger exists.

ID 208

82. Do you take any of these medications regularly to treat **chronic** immune symptoms? (I.e., not to treat acute infections, acute pain, or for cardiovascular health.) *

- ☐ H1 Antihistamines (e.g., Zyrtec, Allegra, Benadryl)
- ☐ H2 Antihistamines (e.g., Pepcid, Zantac, Axid)
- ☐ Leukotriene inhibitors (e.g., Singulair, Accolate, Onon, ZYFLO)
- ☐ Mast Cell Stabilizers (e.g., Ketotifen, Cromolyn, Nedocromil)
- ☐ Non-steroidal Anti-inflammatory Drugs (NSAID) (e.g., aspirin)
- ☐ Inhaled corticosteroids (e.g., Flovent, Pulmicort, Qvar)
- ☐ Bronchodilators (e.g., Albuterol, Levalbuterol, Salmeterol)
- ☐ Expectorants (e.g., Muccinex aka Guaifenisin, Ribavirin, Zanamivir)
- ☐ I don't take any of the above medications on a regular basis

LOGIC Hidden unless: #82 Question "Do you take any of these medications regularly to treat **chronic** immune symptoms? (I.e., not to treat acute infections, acute pain, or for cardiovascular health.)" ("H1 Antihistamines (e.g., Zyrtec, Allegra, Benadryl)","H2 Antihistamines (e.g., Pepcid, Zantac, Axid)")

ID 45

83. Do you take any antihistamines intermitantly, such as seasonally or on an as-needed basis? *

- ○ Yes
- ○ No
- ○ Don't know
- ○ Other, please specify: ______ *

LOGIC Hidden unless: #82 Question "Do you take any of these medications regularly to treat **chronic** immune symptoms? (I.e., not to treat acute infections, acute pain, or for cardiovascular health.)" is one of the following answers ("H1 Antihistamines (e.g., Zyrtec, Allegra, Benadryl)","H2 Antihistamines (e.g., Pepcid, Zantac, Axid)")

ID 186

84. Do you take higher doses of antihistamines than those recommended from the manufacturers? *

- ○ Yes
- ○ Sometimes
- ○ No

LOGIC Hidden unless: (#82 Question "Do you take any of these medications regularly to treat **chronic** immune symptoms? (I.e., not to treat acute infections, acute pain, or for cardiovascular health.)" is one of the following answers ("H1 Antihistamines (e.g., Zyrtec, Allegra, Benadryl)","H2 Antihistamines (e.g., Pepcid, Zantac, Axid)","Leukotriene inhibitors (e.g., Singulair, Accolate, Onon, ZYFLO)","Mast Cell Stabilizers (e.g., Ketotifen, Cromolyn, Nedocromil)") AND #44 Question "Please indicate which, if any, of these types of chronic pain you experience that is unusual for your age (or at one time was) and isn't the result of an acute injury (e.g., a car accident). Please indicate to the right the approximate age around which you started experiencing this pain." is one of the following answers ("Joint pain (arthralgia):","Muscle pain (myalgia):","Arthritis:"))

ID 147

85. Have you ever noticed that your joint/muscle/arthritis pain decreases while taking antihistamines, leukotriene inhibitors, or mast cell stabilizers (or flares up if you go off them)? *

- ○ Yes
- ○ No
- ○ Don't know
- ○ Other, please specify: ________ *

#### Hormones

LOGIC Hidden unless: #3 Question "Sex:" is one of the following answers ("Female")

ID 52

86. Do you have a diagnosis of Polycystic Ovary Syndrome (PCOS)? *

- Yes
- No
- Don't know
- Other, please explain: *

LOGIC Hidden unless: #3 Question "Sex:" is one of the following answers ("Female")

ID 53

87. Are you on any kind of hormone treatment (e.g., birth control) that can potentially affect your menstrual cycle? If so, what type of treatment? *

- Yes: *
- No
- Don't know
- Other, please explain: *

LOGIC Hidden unless: #3 Question "Sex:" is one of the following answers ("Female")

ID 54

88. Do you have or have you ever had any of the following? If you don't experience any of these, please click, "None of the above." (NOTE: Changes to the menstrual cycle should not be the result of conditions such as pregnancy, anorexia, hormone treatment, or similar circumstances. If you have gone through menopause or have had reproductive surgery, e.g., hysterechtomy, please answer about reproductive issues prior to that time period.) *

- [ ] Amenorrhea (one or more missed menstrual periods)
- [ ] Diabetes II or insulin resistance (pre-diabetes)
- [ ] Endometriosis
- [ ] Excessive adult acne
- [ ] Female fertility problems.
- [ ] Frequent dysmenorrhea (severe pain during menstruation)
- [ ] Unusually heavy menstrual bleeding (menorrhagia)
- [ ] High LDL (bad) cholesterol
- [ ] Hypertension (high blood pressure)
- [ ] Hirsutism (excessive hair growth, e.g., face, legs, arms)
- [ ] More frequent menstruation (e.g., every two weeks)
- [ ] Overweight/obese (current or previous)
- [ ] Painful sexual intercourse
- [ ] Premenstrual Dysphoric Disorder (PMDD) - frequent clinically severe PMS
- [ ] Prolonged menstrual bleeding (i.e., more than 5 consecutive days)
- [ ] Severe acne during puberty
- [ ] Uterine fibroids
- [ ] Vaginal dryness
- [ ] None of the above

LOGIC Hidden unless: #3 Question "Sex:" is one of the following answers ("Female")

ID 168

89. Are you going through or have you gone through menopause? *

- Yes
- No
- Not sure

LOGIC Hidden unless: #3 Question "Sex:" is one of the following answers ("Male")

ID 98

90. Do you have or have you ever had any of the following? If you've not experienced any of these things, please check, "None of the above." *

- [ ] Diabetes II or insulin resistance (pre-diabetes)
- [ ] Erectile dysfunction
- [ ] Excessive adult acne
- [ ] Male fertility problems
- [ ] High LDL (bad) cholesterol
- [ ] Hypertension (high blood pressure)
- [ ] Low libido
- [ ] Overweight/obese (current or previous)
- [ ] Severe acne during puberty
- [ ] None of the above

LOGIC Hidden unless: #3 Question "Sex:" is one of the following answers ("Female")

ID 120

91. Have you had a total or partial hysterectomy? *

- Yes, after age 40
- Yes, before age 40
- No

LOGIC Hidden unless: (#90 Question "Do you have or have you ever had any of the following? If you've not experienced any of these things, please check, "None of the above."" is one of the following answers ("Hypertension (high blood pressure)") AND #88 Question "Do you have or have you ever had any of the following? If you don't experience any of these, please click, "None of the above." (NOTE: Changes to the menstrual cycle should not be the result of conditions such as pregnancy, anorexia, hormone treatment, or similar circumstances. If you have gone through menopause or have had reproductive surgery, e.g., hysterechtomy, please answer about reproductive issues prior to that time period.)" is one of the following answers ("Hypertension (high blood pressure)"))

ID 178

92. Are you on any medications to treat your hypertension or other medications that may affect your blood pressure? *

- Yes
- No
- Don't know

LOGIC Show/hide trigger exists. Hidden unless: #3 Question "Sex:" is one of the following answers ("Female")

ID 72

93. Have you ever been pregnant? *

- Yes
- No
- Don't know
- Other, please explain:

*

LOGIC Show/hide trigger exists. Hidden unless: #93 Question "Have you ever been pregnant?" is one of the following answers ("Yes")

ID 73

94. Has at least one of those pregnancies made it to the 3rd trimester? *

- Yes
- No
- Don't know
- Other, please explain:

*

LOGIC Hidden unless: (#93 Question "Have you ever been pregnant?" is one of the following answers ("Yes") AND #94 Question "Has at least one of those pregnancies made it to the 3rd trimester?" is one of the following answers ("No"))

ID 199

95. Did you lose the baby due to miscarriage or some other medical complication? *

- Yes
- No
- Don't know
- Other, please explain: *

LOGIC Hidden unless: #94 Question "Has at least one of those pregnancies made it to the 3rd trimester?" is one of the following answers ("Yes")

ID 74

96. Were any of your children born premature? *

- Yes
- No
- Don't know
- Other, please explain: *

LOGIC Hidden unless: #94 Question "Has at least one of those pregnancies made it to the 3rd trimester?" is one of the following answers ("Yes")

ID 75

97. During any of your pregnancies did you experience preeclampsia (high blood pressure related to pregnancy)? *

- Yes
- No
- Don't know
- Other, please explain: *

LOGIC Hidden unless: #94 Question "Has at least one of those pregnancies made it to the 3rd trimester?" is one of the following answers ("Yes")

ID 76

98. During any of your pregnancies, did you experience gestational diabetes (diabetes arising specifically during pregnancy)? (Please do not answer "yes" if your diabetes began prior to pregnancy.) *

- Yes
- No
- Don't know
- Other, please explain: *

LOGIC Hidden unless: #3 Question "Sex:" is one of the following answers ("Female")

ID 77

99. Have you ever experienced uterine prolapse (collapsed uterus)? *

- Yes
- No
- Don't know
- Other, please explain: *

ID 78

100. Is there anything else about hormone issues you experience that you'd like to tell us about?

#### Family History

ID 79

101. Have you ever had cancer (i.e., a malignancy)? If so, what type? (If you're not certain, please describe it to the best of your abilities.) *

- Yes: *
- No
- Don't know
- Other, please specify: *

LOGIC Show/hide trigger exists.

ID 80

102. Do you know some of the medical history of your biological parents? *

- Yes
- No

LOGIC Hidden unless: #102 Question "Do you know some of the medical history of your biological parents?" is one of the following answers ("Yes")

ID 81

103. Has your mother ever had cancer? If so, what type(s)? *

- Yes:
- No
- Don't know
- Other, please explain: *

LOGIC Hidden unless: #102 Question "Do you know some of the medical history of your biological parents?" is one of the following answers ("Yes")

ID 180

104. Has your mother had a neurodegenerative disorder, such as Alzheimer's Disease, Frontotemporal Dementia, or Parkinson's? If so, what type(s)? *

- Yes:
- Maybe
- No
- Other, please explain: *

LOGIC Hidden unless: #102 Question "Do you know some of the medical history of your biological parents?" is one of the following answers ("Yes")

ID 82

105. Has your father ever had cancer? If so, what type(s)? *

- Yes: *
- No
- Don't know
- Other, please explain: *

LOGIC Hidden unless: #102 Question "Do you know some of the medical history of your biological parents?" is one of the following answers ("Yes")

ID 181

106. Has your father had a neurodegenerative disorder, such as Alzheimer's Disease, Frontotemporal Dementia, or Parkinson's? If so, what type(s)? *

- Yes:
- Maybe
- No
- Other, please explain:

*

LOGIC Show/hide trigger exists.

ID 182

107. Do you know some of the medical history of your biological grandparents? *

- Yes
- No

LOGIC Hidden unless: #107 Question "Do you know some of the medical history of your biological grandparents?" is one of the following answers ("Yes")

ID 183

108. Please check which of your grandparents has had cancer and in the box provided list the type(s) of cancer. If you're not certain, answer to the best of your abilities. *

- [ ] Paternal grandfather: ________ *
- [ ] Paternal grandmother: ________ *
- [ ] Maternal grandfather: ________ *
- [ ] Maternal grandmother: ________ *
- [ ] None of the above/Don't know

LOGIC Hidden unless: #107 Question "Do you know some of the medical history of your biological grandparents?" is one of the following answers ("Yes")

ID 184

109. Please check which of your grandparents has had a neurodegenerative disorder and in the box provided list the type(s) of conditions. If you're not certain, answer to the best of your abilities *

- [ ] Paternal grandfather: ________ *
- [ ] Paternal grandmother: ________ *
- [ ] Maternal grandfather: ________ *
- [ ] Maternal grandmother: ________ *
- [ ] None of the above/Don't know

#### Anything else?

ID 192

110. Do you (or your adult ward) have any other medical issues you'd like to tell us about that haven't been covered in this survey?

#### Future Research

LOGIC Hidden unless: (#12 Question "Do you have a professional diagnosis of Autism Spectrum Disorder (ASD) or a related diagnosis (e.g., Asperger's Syndrome)? If so, please write the diagnosis in the box provided." is one of the following answers ("Yes:") OR #18 Question "What type of EDS are you diagnosed with? If you are not diagnosed with EDS but with Joint Hypermobility Syndrome (JHS), please click the option below." is one of the following answers ("EDS, Hypermobility Type"))

ID 136

111. Have you had informal genotyping performed through commercial sites such as 23andme.com or Ancestry.com? If so, please list the site(s) you've used in the space provided. *

- Yes: *
- No

LOGIC Hidden unless: (((#12 Question "Do you have a professional diagnosis of Autism Spectrum Disorder (ASD) or a related diagnosis (e.g., Asperger's Syndrome)? If so, please write the diagnosis in the box provided." is one of the following answers ("No") AND #14 Question "Do any of your 1st degree **biological** relatives (parent, sibling, child) have an ASD diagnosis?" is one of the following answers ("No")) AND #17 Question "Do you have a professional diagnosis of Ehlers-Danlos Syndrome (EDS) or Joint Hypermobility Syndrome (JHS)?" is one of the following answers ("No")) AND #23 Question "Do any of your 1st degree **biological** relatives (parent, sibling, child) have a diagnosis of EDS or JHS? Please check all that apply." is one of the following answers ("No"))

ID 204

112. Have you had informal genotyping performed through commercial sites such as 23andme.com or Ancestry.com? If so, please list the site(s) you've used in the space provided. *

- Yes: *
- No

LOGIC Show/hide trigger exists. Hidden unless: (#111 Question "Have you had informal genotyping performed through commercial sites such as 23andme.com or Ancestry.com? If so, please list the site(s) you've used in the space provided." is one of the following answers ("Yes:") OR #112 Question "Have you had informal genotyping performed through commercial sites such as 23andme.com or Ancestry.com? If so, please list the site(s) you've used in the space provided." is one of the following answers ("Yes:"))

ID 137

113. Would you be interested in providing your raw genetic data for future research into genetic variations that could be involved in risk for ASD and H-EDS/JHS? (Please be aware, you are not currently being asked to provide your raw data for this survey, nor is your answer to this question a promise to participate. We are simply trying to gauge future interest.) *

- ○ Yes, I would be interested in providing my raw data for genetics research into ASD and H-EDS/JHS
- ○ Maybe
- ○ No, I'm not interested

LOGIC Hidden unless: #113 Question "Would you be interested in providing your raw genetic data for future research into genetic variations that could be involved in risk for ASD and H-EDS/JHS? (Please be aware, you are not currently being asked to provide your raw data for this survey, nor is your answer to this question a promise to participate. We are simply trying to gauge future interest.)" is one of the following answers ("Yes, I would be interested in providing my raw data for genetics research into ASD and H-EDS/JHS")

ID 198

114. If you would like us to get back in touch with you for future genetics studies, please provide your email address in the box below. Your email address will be kept strictly confidential and will be stored in a secure computer system. It will not be used for anything other than the studies mentioned here.

**Thank You!**

ID 1

Thank you! If you have any questions, comments, or concerns, please feel free to contact Dr. Emily Casanova at or Dr. Manuel Casanova at.
