## Supplementary File - Statistics & Additional Results for "Immune, Autonomic, and Endocrine Dysregulation in Autism and Ehlers-Danlos Syndrome/Hypermobility Spectrum Disorders Versus Unaffected Controls"

**Title:** A Cohort Study Comparing Medical Issues in Ehlers-Danlos Syndrome/Hypermobility Spectrum Disorders, Autism Spectrum Disorder, and Unaffected Controls.

**Contact:** Emily L. Casanova at

Dept. Biomedical Sciences, University of South Carolina School of Medicine Greenville, Greenville, South Carolina, USA.

**Supplementary File 2**

**Supplementary Table 1.** Statistical results of between-group and within-group sex differences in immune-mediated, endocrine-mediated, and autonomic symptoms.

| **Between-Group and Sex Symptom Differences** | |  |  |  |
| --- | --- | --- | --- | --- |
|  | **Between-Group Differences (Males)** | | **Between-Group Differences (Females)** | **Within-Group Sex Differences** |
| **Immune-mediated Symptoms** | *p* = 0.003*  *X^2^* = 11.782 | | *p* < 0.0001*  *X^2^* = 135.64 | **EDS/HSD:**  *p* < 0.001*  *W* = 3081  **Autism:**  *p* = 0.005*  *W* = 3018  **CON:**  *p* = 0.024*  *W* = 2826.5 |
| **Endocrine-mediated Symptoms** | *p* = 0.4302  *X^2^* = 1.6871 | | *p* < 0.0001*  *X^2^* = 55.792 | **EDS/HSD:**  *p* = 0.562  *W* = 2149  **Autism:**  *p* = 0.913  *W* = 2333  **CON:**  *p* = 0.748  *W* = 2373  (abbreviated symptom list) |
| **Autonomic Symptoms** | *p* < 0.0001*  *X^2^ =* 26.60 | | *p* < 0.0001*  *X^2^* = 217.57 | **EDS/HSD:**  *p* = 0.003*  *W* = 2856  **Autism:**  *p* = 0.016*  *W* = 2926.5  **CON:**  *p* = 0.021*  *W* = 2843 |

**Supplementary Table 2.** Immune-mediated, endocrine-mediated, and autonomic nervous system symptoms by EDS/HSD subgroup.

| **Symptoms by Group** |  |  |  |  |  |  |  |
| --- | --- | --- | --- | --- | --- | --- | --- |
| **System** | **Symptom** | **hEDS (N = 214)** | **cEDS (N = 27)** | **vEDS (N = 9)** | **JHS (N = 26)** | **Autism (N = 89)** | **CON (N = 98)** |
| **Immune** | *Mast Cell Activation Syndrome (MCAS)* | 7% | 4% | 0% | 4% | 1% | 0% |
|  | *Anaphylaxis* | 21% | 30% | 11% | 15% | 3% | 1% |
|  | *Respiratory Allergies* | 50% | 48% | 33% | 46% | 31% | 20% |
|  | *Asthma* | 39% | 41% | 11% | 35% | 24% | 13% |
|  | *Childhood Ear Infections* | 43% | 59% | 22% | 27% | 40% | 30% |
|  | *Adult Ear Infections* | 16% | 37% | 22% | 19% | 8% | 9% |
|  | *Chronic Rhinitis* | 34% | 44% | 0% | 31% | 20% | 8% |
|  | *Chronic Sinusitis* | 51% | 63% | 22% | 38% | 31% | 18% |
|  | *Dry, Itchy Eyes* | 52% | 44% | 33% | 42% | 25% | 12% |
|  | *Skin Allergies* | 50% | 70% | 11% | 38% | 30% | 14% |
|  | *Eczema* | 33% | 26% | 11% | 42% | 26% | 11% |
|  | *Hives, Rash* | 48% | 63% | 33% | 38% | 25% | 8% |
|  | *Food Allergies* | 38% | 48% | 44% | 31% | 29% | 10% |
|  | *Unusual Physical Reactions to Medications* | 50% | 70% | 44% | 42% | 33% | 9% |
|  | *Unusual Physical Reactions to Environmental Chemicals* | 51% | 52% | 67% | 35% | 22% | 10% |
|  | *Unusual Physical Reactions to Stress* | 39% | 33% | 22% | 31% | 44% | 10% |
|  | *Recurrent Urinary Tract Infections or Inflammation* | 37% | 37% | 67% | 23% | 16% | 11% |
|  | *Recurrent Vaginal Infections or Inflammation* | 25% | 19% | 44% | 31% | 11% | 4% |
|  | *Stomach Ulcers* | 16% | 19% | 33% | 0% | 7% | 3% |
|  | *Acid Reflux* | 59% | 56% | 56% | 31% | 24% | 18% |
|  | *Transiently Swollen Lymph Nodes* | 34% | 37% | 22% | 8% | 16% | 3% |
|  | *Autoimmune Disorder (self)* | 17% | 19% | 11% | 15% | 11% | 12% |
|  | *Autoimmune Disorder (1^st^ Degree Relative)* | 30% | 44% | 22% | 38% | 20% | 12% |
|  | *Adverse Reaction to Vaccination* | 29% | 41% | 11% | 12% | 18% | 6% |
|  | *Difficulty Developing Immunity Following Vaccination or Illness* | 25% | 33% | 33% | 12% | 9% | 8% |
|  | *Avg. # of Immune Symptoms Per Person* | 8 | 9.1 | 6.2 | 6.2 | 4.8 | 2.4 |
| **System** | **Symptom** | **hEDS (N = 214)** | **cEDS (N = 27)** | **vEDS (N = 9)** | **JHS (N = 26)** | **Autism (N = 89)** | **CON (N = 98)** |
| **Endocrine** | *Polycystic Ovary Syndrome (PCOS)* | 17% | 26% | 22% | 4% | 8% | 6% |
|  | *Amenorrhea* | 33% | 41% | 11% | 27% | 24% | 32% |
|  | *Diabetes II or Prediabetes* | 7% | 11% | 11% | 12% | 11% | 4% |
|  | *Endometriosis* | 22% | 33% | 44% | 8% | 17% | 2% |
|  | *Adult Acne* | 22% | 15% | 0% | 8% | 22% | 11% |
|  | *Female Fertility Issues* | 17% | 15% | 22% | 12% | 7% | 8% |
|  | *Dysmenorrhea* | 61% | 63% | 67% | 31% | 40% | 20% |
|  | *Heavy Menses* | 58% | 74% | 67% | 42% | 33% | 20% |
|  | *High LDL Cholesterol* | 15% | 22% | 22% | 12% | 11% | 7% |
|  | *Hypertension* | 11% | 26% | 11% | 19% | 6% | 14% |
|  | *Hirsutism* | 15% | 22% | 0% | 12% | 19% | 7% |
|  | *Frequent Menstruation* | 22% | 26% | 44% | 15% | 16% | 8% |
|  | *Overweight* | 42% | 48% | 67% | 50% | 44% | 34% |
|  | *Painful Sex* | 45% | 30% | 22% | 38% | 21% | 13% |
|  | *Premenstrual Dysphoric Disorder (PMDD)* | 20% | 7% | 11% | 4% | 16% | 4% |
|  | *Prolonged Menstruation* | 49% | 63% | 78% | 35% | 38% | 20% |
|  | *Severe Acne in Puberty* | 14% | 26% | 0% | 12% | 26% | 16% |
|  | *Uterine Fibroids* | 14% | 19% | 22% | 12% | 10% | 5% |
|  | *Vaginal Dryness* | 29% | 22% | 33% | 15% | 11% | 14% |
|  | *Avg. # of Endocrine Symptoms Per Person* | 5.1 | 5.9 | 5.6 | 3.7 | 3.8 | 2.5 |
| **System** | **Symptom** | **hEDS (N = 214)** | **cEDS (N = 27)** | **vEDS (N = 9)** | **JHS (N = 26)** | **Autism (N = 89)** | **CON (N = 98)** |
| **Autonomic** | *Postural Orthostatic Tachycardia Syndrome (POTS)* | 29% | 30% | 63% | 4% | 1% | 0% |
|  | *Chronic Fatigue* | 90% | 78% | 78% | 73% | 54% | 29% |
|  | *Dizziness, Vertigo* | 91% | 81% | 100% | 81% | 55% | 29% |
|  | *Fainting* | 28% | 37% | 33% | 31% | 4% | 2% |
|  | *Brain Fog* | 83% | 70% | 89% | 81% | 46% | 26% |
|  | *Heart Palpitations* | 76% | 63% | 67% | 46% | 29% | 17% |
|  | *Chest Pains* | 45% | 33% | 44% | 23% | 13% | 5% |
|  | *Shortness of Breath* | 59% | 56% | 44% | 42% | 22% | 8% |
|  | *Weakness* | 70% | 59% | 67% | 42% | 28% | 15% |
|  | *Non-migraine Headaches* | 66% | 74% | 67% | 46% | 35% | 24% |
|  | *Abnormal Sweating* | 65% | 56% | 78% | 46% | 31% | 14% |
|  | *Gastrointestinal Distress* | 83% | 70% | 89% | 69% | 55% | 20% |
|  | *Muscle Tremors* | 56% | 56% | 56% | 42% | 18% | 5% |
|  | *Bladder Dysfunction* | 42% | 52% | 22% | 23% | 18% | 3% |
|  | *Insomnia* | 68% | 89% | 67% | 50% | 49% | 31% |
|  | *Free-floating Anxiety* | 67% | 67% | 33% | 69% | 58% | 38% |
|  | *Unusual Blood Pressure Fluctuations* | 41% | 48% | 44% | 42% | 9% | 7% |
|  | *Unusually High or Low Body Temperature* | 86% | 85% | 89% | 77% | 47% | 23% |
|  | *Avg. # of Autonomic Symptoms Per Person* | 11.2 | 10.7 | 10.8 | 8.8 | 5.7 | 3 |

**Supplementary Table 3.** Immune-mediated, endocrine-mediated, and autonomic symptoms by EDS/HSD subgroup.

| **Symptoms by Group** |  |  |  |  |  |  |  |
| --- | --- | --- | --- | --- | --- | --- | --- |
| **System** | **Symptom** | **EDS/HSD (f) (*N* = 248)** | **EDS/HSD (m) (*N* = 16)** | **Autism (f) (*N* = 89)** | **Autism (m) (*N* = 53)** | **CON (f) (*N* = 98)** | **CON (m) (*N* = 47)** |
| **Immune** | *Anaphylaxis* | 19% | 19% | 3% | 4% | 1% | 0% |
|  | *Respiratory Allergies* | 48% | 25% | 31% | 15% | 20% | 11% |
|  | *Asthma* | 37% | 31% | 24% | 25% | 13% | 15% |
|  | *Childhood Ear Infections* | 43% | 25% | 40% | 23% | 30% | 21% |
|  | *Adult Ear Infections* | 18% | 0% | 8% | 2% | 9% | 0% |
|  | *Chronic Rhinitis* | 32% | 19% | 20% | 17% | 8% | 6% |
|  | *Chronic Sinusitis* | 51% | 31% | 31% | 19% | 18% | 6% |
|  | *Dry, Itchy Eyes* | 49% | 25% | 25% | 17% | 12% | 11% |
|  | *Skin Allergies* | 46% | 19% | 30% | 11% | 14% | 6% |
|  | *Eczema* | 29% | 0% | 26% | 23% | 11% | 15% |
|  | *Hives, Rash* | 46% | 13% | 25% | 9% | 8% | 2% |
|  | *Food Allergies* | 35% | 19% | 29% | 11% | 10% | 9% |
|  | *Unusual Reactions to Medications* | 50% | 25% | 33% | 13% | 9% | 4% |
|  | *Unusual Reactions to Environmental Chemicals* | 48% | 25% | 22% | 9% | 10% | 6% |
|  | *Unusual Reactions to Stress* | 37% | 25% | 44% | 30% | 10% | 4% |
|  | *Urinary Tract Infection or Inflammation* | 39% | 0% | 16% | 4% | 11% | 0% |
|  | *Stomach Ulcers* | 14% | 13% | 7% | 9% | 3% | 2% |
|  | *Acid Reflux* | 55% | 38% | 24% | 34% | 18% | 26% |
|  | *Transiently Swollen Lymph Nodes* | 32% | 0% | 16% | 4% | 3% | 2% |
|  | *Autoimmune Disorders* | 18% | 13% | 11% | 9% | 12% | 2% |
|  | *Avg. # of Immune Symptoms Per Person* | 7.4 | 3.6 | 4.7 | 2.9 | 2.3 | 1.5 |
| **System** | **Symptom** | EDS/HSD (f) (*N* = 248) | EDS/HSD (m) (*N* = 16) | Autism (f) (*N* = 89) | Autism (m) (*N* = 53) | CON (f) (*N* = 98) | CON (m) (*N* = 47) |
| **Endocrine** | *Diabetes II or Prediabetes* | 8% | 13% | 11% | 8% | 4% | 0% |
|  | *Adult Acne* | 21% | 6% | 22% | 19% | 11% | 6% |
|  | *High LDL Cholesterol* | 18% | 31% | 11% | 23% | 7% | 11% |
|  | *Hypertension* | 14% | 6% | 6% | 23% | 14% | 11% |
|  | *Overweight* | 44% | 25% | 44% | 36% | 33% | 40% |
|  | *Severe Acne in Puberty* | 14% | 25% | 26% | 15% | 16% | 17% |
|  | *Amennorrhea* | 35% | N/A | 24% | N/A | 32% | N/A |
|  | *Endometriosis* | 24% | N/A | 17% | N/A | 2% | N/A |
|  | *Female Fertility Issues* | 16% | N/A | 7% | N/A | 8% | N/A |
|  | *Dysmenorrhea* | 60% | N/A | 40% | N/A | 20% | N/A |
|  | *Heavy Menstruation* | 58% | N/A | 33% | N/A | 20% | N/A |
|  | *Hirsutism* | 17% | N/A | 19% | N/A | 7% | N/A |
|  | *Frequent Menstruation* | 24% | N/A | 16% | N/A | 8% | N/A |
|  | *Painful Sex* | 44% | N/A | 21% | N/A | 13% | N/A |
|  | *Polycystic Ovary Syndrome* | 17% | N/A | 8% | N/A | 6% | N/A |
|  | *Premenstrual Dysphoric Disorder* | 20% | N/A | 16% | N/A | 4% | N/A |
|  | *Prolonged Menstruation* | 52% | N/A | 38% | N/A | 20% | N/A |
|  | *Uterine Fibroids* | 16% | N/A | 10% | N/A | 5% | N/A |
|  | *Vaginal Dryness* | 28% | N/A | 11% | N/A | 14% | N/A |
|  | *Avg. # of Endocrine Symptoms Per Person* | 5.1 | 1 | 3.8 | 1.2 | 2.5 | 0.9 |
| **System** | **Symptom** | EDS/HSD (f) (*N* = 248) | EDS/HSD (m) (*N* = 16) | Autism (f) (*N* = 89) | Autism (m) (*N* = 53) | CON (f) (*N* = 98) | CON (m) (*N* = 47) |
| **Autonomic** | *Postural Orthostatic Tachycardia Syndrome* | 29% | 13% | 1% | 0% | 0% | 0% |
|  | *Chronic Fatigue* | 88% | 94% | 53% | 42% | 29% | 15% |
|  | *Dizziness, Vertigo* | 89% | 63% | 55% | 36% | 29% | 9% |
|  | *Fainting* | 29% | 0% | 4% | 0% | 2% | 0% |
|  | *Brain Fog* | 83% | 69% | 46% | 38% | 26% | 11% |
|  | *Heart Palpitations* | 75% | 38% | 29% | 17% | 17% | 13% |
|  | *Chest Pains* | 42% | 25% | 13% | 13% | 5% | 4% |
|  | *Shortness of Breath* | 58% | 31% | 22% | 15% | 8% | 9% |
|  | *Weakness* | 68% | 50% | 28% | 23% | 15% | 6% |
|  | *Non-migraine Headaches* | 67% | 56% | 35% | 28% | 24% | 17% |
|  | *Abnormal Sweating* | 64% | 50% | 31% | 34% | 14% | 19% |
|  | *Gastrointestinal Distress* | 81% | 56% | 55% | 25% | 20% | 23% |
|  | *Muscle Tremors* | 55% | 25% | 18% | 17% | 5% | 11% |
|  | *Bladder Dysfunction* | 39% | 19% | 18% | 11% | 3% | 2% |
|  | *Insomnia* | 71% | 63% | 49% | 40% | 31% | 21% |
|  | *Free-floating Anxiety* | 65% | 63% | 58% | 47% | 38% | 21% |
|  | *Unusual Blood Pressure Fluctuations* | 42% | 31% | 9% | 9% | 7% | 2% |
|  | *Unusually High or Low Body Temperature* | 85% | 38% | 47% | 30% | 23% | 11% |
|  | *Avg. # of Autonomic Symptoms Per Person* | 11 | 7.7 | 5.7 | 4.3 | 3 | 1.9 |

**Supplementary Figure 1.** Correlations between total number of symptoms across the immune, endocrine, and autonomic systems reviewed in this study. The more symptoms reported in relation to a single system, the more likely the individual will experience increased symptoms within the other two systems.

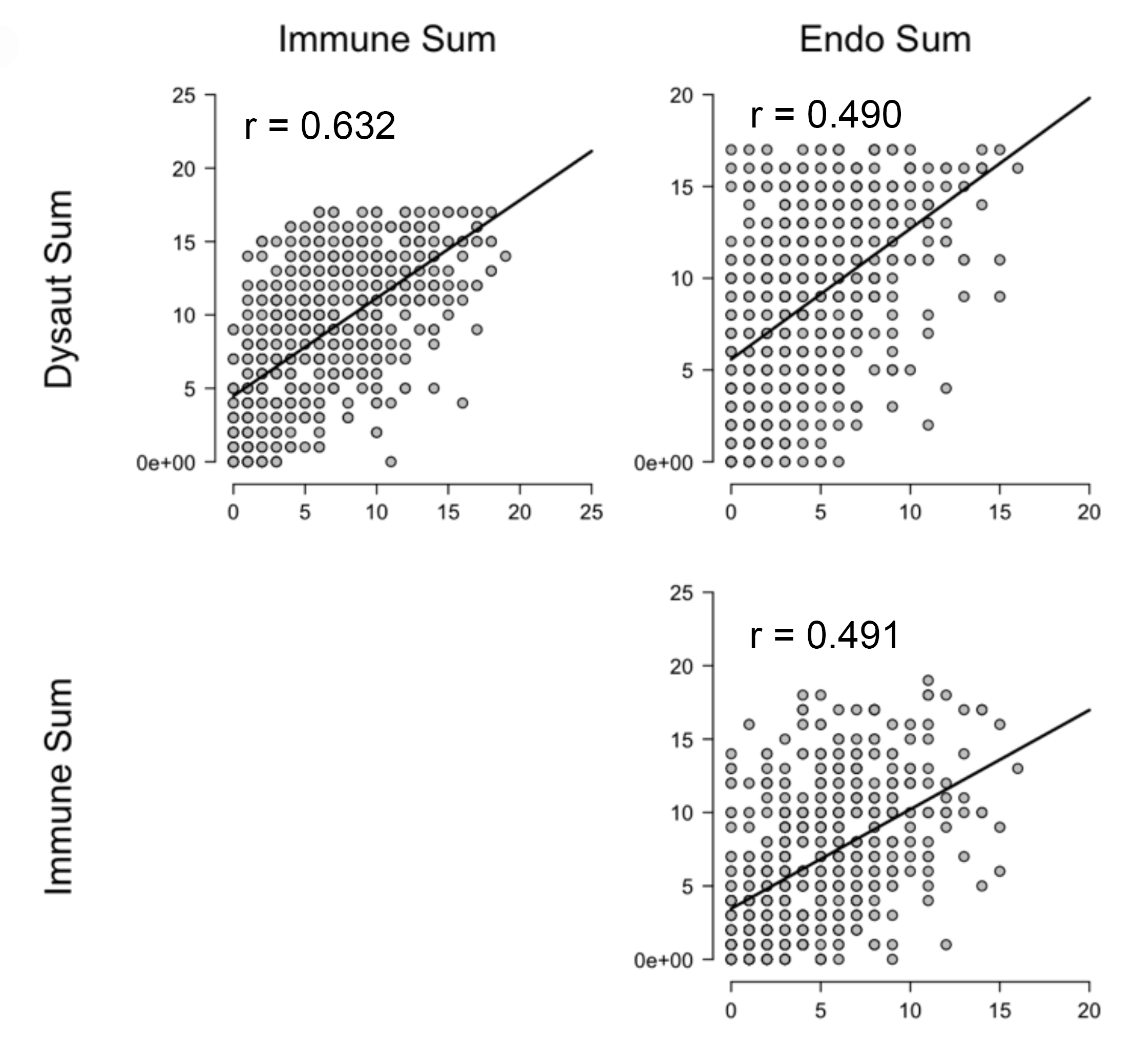

**Further Data on Immune Dysregulation**

The log odds significantly differed across groups concerning adverse reactions to vaccinations and challenges developing proper immunity to inoculation or wild-type illness (e.g., presenting with negative titers following vaccination or contracting an illness such as chicken pox on multiple occasions) [each *p* < 0.0001] (see Supplementary File 2, Fig. 2). EDS/HSD women reported the highest rates of adverse vaccination reaction (28%) and mild immunodeficiencies (25%) compared to autism (18%, 9%, respectively) and controls (6%, 8%, respectively). Of those individuals reporting adverse reaction following vaccination, irrespective of clinical group, the majority (57%) reportedly reacted to the influenza vaccine (Supplementary File 2, Figure 3). Meanwhile, the odds of mild immunodeficiencies for EDS/HSD women is about 3 times higher than the odds for autistic women (95% CI: 1.20, 9.12) and about 3.7 times higher than the odds for control women (95% CI: 1.33, 10.10). Anecdotally, these traits are common in the condition known as mast cell activation syndrome (MCAS), which is frequently comorbid with EDS/HSD and has also previously been noted in autism

**Supplementary Figure 2.** (A) Individuals reporting adverse reaction to vaccination broken down according to clinical group. (B) Individuals reporting mild immunodeficiencies broken down according to clinical group.

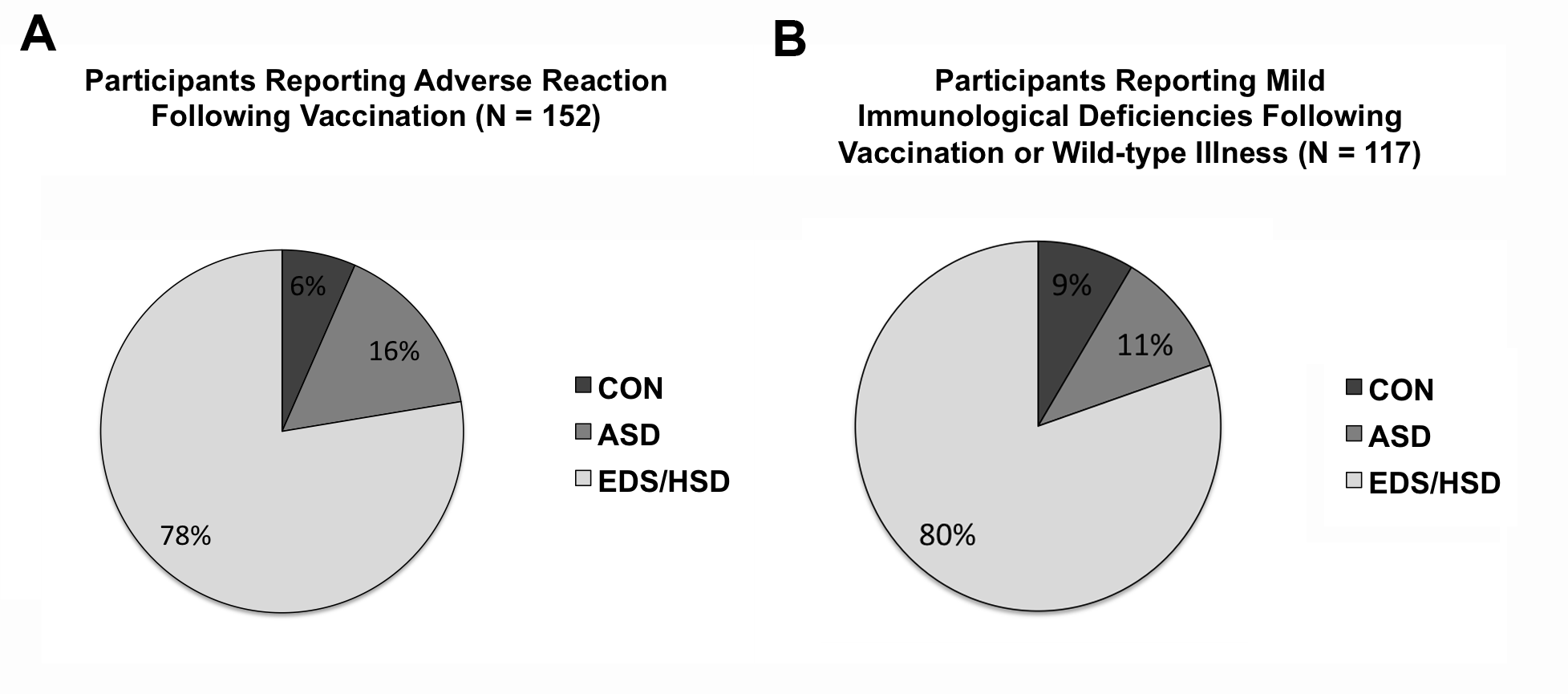
**Supplementary Figure 3.** Breakdown of specific adverse vaccination reactions reported by respondents. As the histogram illustrates, influenza vaccination reportedly causes the most problems for our sample, regardless of clinical group. However, individuals are also likely to be exposed to the influenza vaccine on a more frequent basis (e.g., yearly) compared to vaccinations such as tetanus, potentially increasing the frequency of exposure and sensitization. Anecdotal evidence of those with EDS/HSD, however, suggests that some individuals may also have serious adverse reactions to wild-type illness, indicating that vaccinations should not be avoided entirely but applied with caution and used on a case-by-case basis.

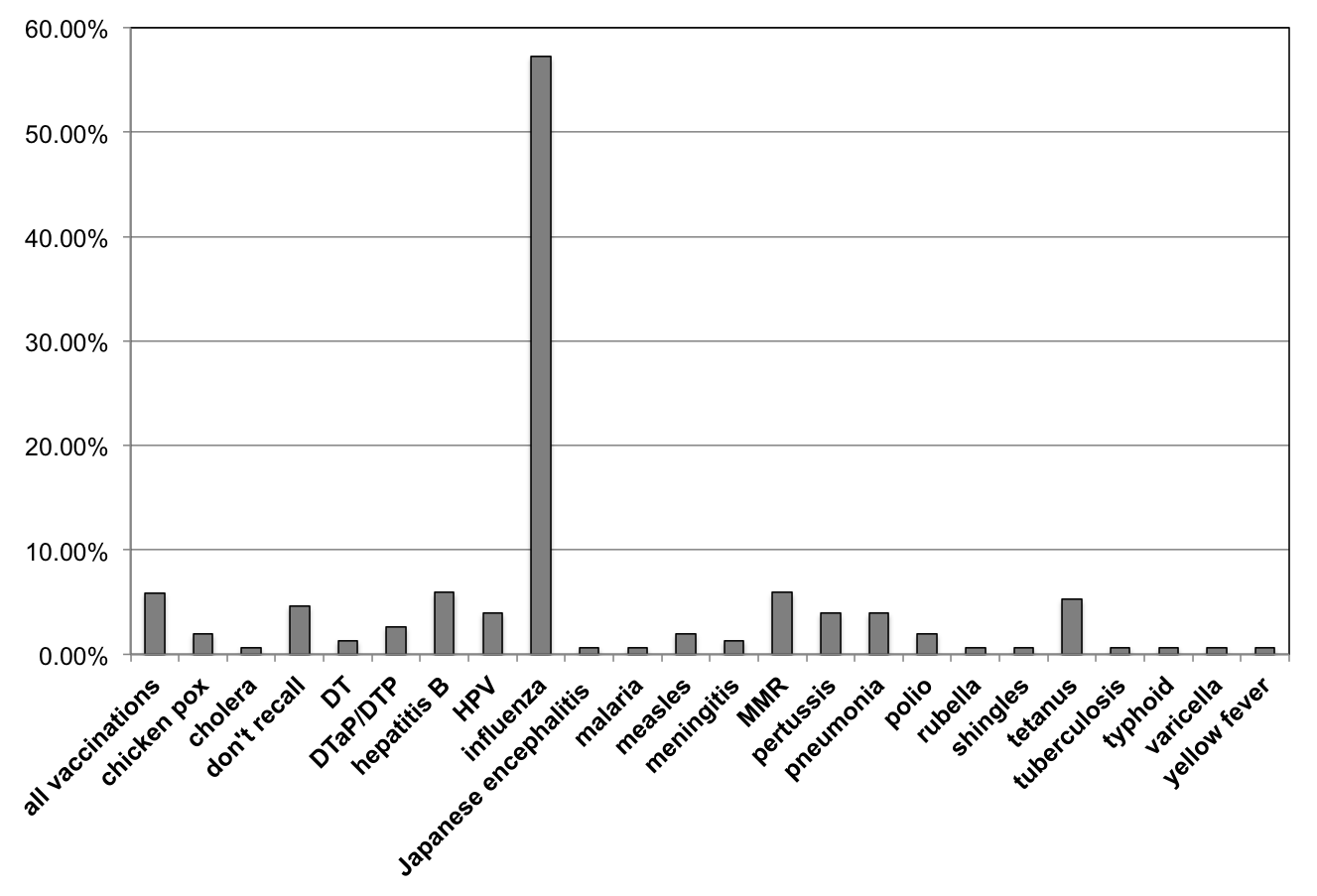

**Gastrointestinal Distress and Its Relationship to Autonomic Symptoms**

While gastrointestinal (GI) distress was an item included within the autonomic panel, we found that individuals reporting chronic GI distress reported more autonomic symptoms in general, irrespective of group, suggesting that the autonomic nervous system may be playing a significant role in GI disorders in both clinical and non-clinical populations [*F* = 335.058, *p* < 0.001].
